# Organotin(IV) Dithiocarbamate Compounds Targeting A549 Lung Cancer Cells via Mitochondria-Mediated Apoptosis

**DOI:** 10.64898/2026.03.26.714399

**Authors:** Nurul Amalina Abd Aziz, Normah Awang, Nurul Farahana Kamaludin, Asmah Hamid, Nur Najmi Mohamad Anuar, Kok Meng Chan, Nurul Zahidah Zainirizal

**Affiliations:** Center for Toxicology and Health Risk Studies, Faculty of Health Sciences, Universiti Kebangsaan Malaysia, Jalan Raja Muda Abdul Aziz, 50300 Kuala Lumpur, Malaysia; Product Stewardship and Toxicology, Petroliam Nasional Berhad, Level 13, Tower 1, PETRONAS Twin Towers, KLCC, 50088 Kuala Lumpur, Malaysia; Environmental Health & Industrial Safety Programme, Faculty of Health Sciences, Universiti Kebangsaan Malaysia, Jalan Raja Muda Abdul Aziz, 50300 Kuala Lumpur, Malaysia

**Keywords:** Lung cancer therapy, Organotin(IV) dithiocarbamates, Apoptosis induction, A549 cell line

## Abstract

Lung cancer remains the leading cause of cancer-related deaths worldwide, with cisplatin as the primary chemotherapy despite its limitations. Organotin(IV) dithiocarbamates have emerged as promising anticancer agents due to their potent cytotoxicity and stability. This study reports the successful synthesis of four novel organotin(IV) dithiocarbamates: dimethyltin(IV) *N*-methyl-*N*-benzyldithiocarbamate (DioSn-**1**), diphenyltin(IV) *N*-methyl-*N*-benzyldithiocarbamate (DioSn-**2**), triphenyltin(IV) *N*-methyl-*N*-benzyldithiocarbamate (TriSn-**3**), and triphenyltin(IV) *N*-ethyl-*N*-benzyldithiocarbamate (TriSn-**4**). Their cytotoxicity against A549 lung carcinoma cells was evaluated via MTT assay, while Annexin V-FITC/PI staining determined the mode of cell death. DioSn-**2**, TriSn-**3**, and TriSn-**4** exhibited potent cytotoxicity (IC₅₀: 0.52–1.86 μM), whereas DioSn-**1** was inactive (IC₅₀ > 50 μM). Apoptotic features such as cell shrinkage and membrane blebbing were observed, with apoptosis rates ranging from 58% to 91%. DioSn-**2** was the most selective (SI = 6.45) and induced early DNA damage within 30 minutes, followed by mitochondrial depolarization and excessive ROS generation. Caspase-9 activation exceeded caspase-8, confirming intrinsic apoptosis. NAC treatment reduced apoptosis by 52%, highlighting oxidative stress as a key cytotoxic mechanism. These findings suggest DioSn-**2** as a promising alternative to cisplatin for lung cancer therapy.

## INTRODUCTION

Lung cancer remains the leading cause of cancer incidence and mortality worldwide, with approximately 2 million cases and 1.8 million deaths reported annually [1,2]. It is classified into two major histological subtypes: small cell lung cancer (SCLC), which accounts for 10–15% of cases [3] and non-small cell lung cancer (NSCLC), which constitutes approximately 80% of cases [4]. Recent data indicate that NSCLC remains the predominant cause of cancer-related morbidity globally [5]. While risk factors for lung cancer vary by region, smoking remains the primary cause worldwide [6], alongside environmental pollution and dietary habits [7].

Current lung cancer treatments include chemotherapy, chemoradiotherapy, targeted therapy, anti-angiogenic therapy, immunotherapy, and combination regimens [8]. Since the 1970s, cisplatin has been the cornerstone of NSCLC chemotherapy, exerting its cytotoxic effects by inducing genotoxicity and promoting the production of reactive oxygen species (ROS), thereby triggering apoptosis [9]. Unlike radiotherapy or surgery, chemotherapy acts systemically to prevent cancer progression and metastasis [10]. The success of cisplatin has driven interest in metal-based anticancer agents, including organometallic and inorganic compounds. However, chemotherapy resistance remains a major challenge, reducing drug efficacy in subsequent treatments. Furthermore, the severe side effects of platinum-based drugs limit their clinical application, underscoring the need for safer and more effective alternatives [11].

Anticancer metallodrugs primarily exert their effects by interacting with DNA, which plays a crucial role in gene expression, transcription, mutagenesis, and protein synthesis. Beyond DNA, these metal complexes also target key cellular proteins involved in antioxidant defense, DNA repair, and electron transport [12,13]. Organotin(IV) compounds have emerged as promising alternatives to platinum-based drugs due to their ability to target DNA while exhibiting higher antiproliferative activity, improved elimination rates, and reduced side effects [14]. Consequently, extensive research has focused on novel organotin(IV) compounds derived from dithiocarbamate ligands to enhance treatment efficacy while minimizing adverse effects [14,15]. Organotin compounds, which consist of tin bound to hydrocarbons [16], possess diverse biological activities influenced by their structural variations [17]. Notably, organotin(IV) dithiocarbamate compounds contain two sulfur atoms in their ligand, which strongly bind to metals and enhance their bioactive potential [18].

These compounds form stable complexes with transition metals, making them valuable in multiple applications, including agriculture, biological research, and catalysis. Additionally, they serve as precursors for tin sulfide nanoparticles [19]. Organotin(IV) dithiocarbamate compounds exhibit significant antitumor and antibacterial properties, highlighting their potential as chemotherapeutic agents [14,20]. These compounds have demonstrated strong antiproliferative effects in vitro and are of growing interest due to their ability to stabilize specific stereochemistry, further enhancing their therapeutic potential [17].

Previous studies have reported the synthesis and cytotoxic activity of certain organotin(IV) dithiocarbamate compounds using the MTT assay [21]. However, their selectivity and apoptotic mechanisms remain largely unexplored. This study aims to address this gap by investigating newly synthesized di- and triorganotin(IV) *N*-alkyl-*N*-benzyldithiocarbamate compounds for their cytotoxic effects on human lung carcinoma (A549) cells. By expanding on both structural characterization and mechanistic evaluation, this research provides deeper insights into their potential as alternative chemotherapeutic agents, overcoming the limitations of conventional platinum-based therapies.

## MATERIALS AND METHODS

### Synthesize and characterization of compounds

*N*-ethylbenzylamine, *N*-methylbenzylamine, and organotin(IV) chlorides were purchased from Sigma Aldrich and Alfa Aesar, while carbon disulfide, ammonium solution, chloroform, and ethanol were obtained from Merck and Systerm Chemicals. All chemicals were used as received, following the suppliers’ guidelines. Characterization techniques included melting point determination (MPA120 EZ-Melt), elemental analysis (PerkinElmer 2400), and infrared spectroscopy (Bruker Vertex 70v FT-IR, 4000–80 cm⁻¹). NMR spectroscopy (^13^C and ^119^Sn) was performed using Bruker Avance III (400 MHz) with CDCl₃ as the solvent and tetramethylsilane as the internal standard. Dithiocarbamate ligands were synthesized via an *in-situ* method under ambient conditions, as described by Kamaludin and Awang (2014). Secondary amines in ethanol were stirred with ammonium solution to create a basic medium, followed by the addition of carbon disulfide under cold conditions (<4 °C) to form the ligands. These ligands were then reacted with organotin(IV) salts in specific molar ratios (2:1 for diorganotin, 1:1 for triorganotin complexes) to synthesize organotin(IV) complexes. The reaction mixtures were stirred continuously, and the resulting precipitates were filtered, washed with cold ethanol, and vacuum-dried to obtain pure solid products. Crystals of DioSn-**1**, TriSn-**3**, and TriSn-**4** were successfully grown via slow evaporation of a chloroform-ethanol solution (1:2; 2:1 v/v) over 3–4 days, while DioSn-**2** failed to crystallize under similar conditions. X-ray crystallography was performed on the obtained crystals, with TriSn-4’s structure previously reported [21], offering valuable structural insights.

### Preparation of Stock Compounds

Stock solutions were prepared at 20 mM for DioSn-**1**, TriSn-**3**, and TriSn-**4**, and 5 mM for DioSn-**2**. Each compound was accurately weighed, dissolved in DMSO, and stored at 4°C. Fresh dilutions were made in culture media before experiments.

### Cell Culture and Reagents

The A549 cell line (ATCC-CCL-185) and MRC-5 cell line (ATCC-CCL-171) were obtained from the American Type Culture Collection (ATCC). A549 cells were cultured in Kaighn’s Modification of Ham’s F-12 Medium (F-12K), while MRC-5 cells were maintained in Eagle’s Minimum Essential Medium (EMEM), both supplemented with 10% fetal bovine serum (FBS) and 1% penicillin-streptomycin. The cells were incubated at 37°C in a humidified atmosphere with 5% CO₂.

### MTT Assay for Cytotoxic Assessment

The MTT assay was performed following Mosmann’s method [22]. A549 cells (5 × 10⁴ cells/mL) were seeded in 96-well plates and treated for 24 hours with an initial highest concentration of 50 μM (DioSn-**1**), 5 μM (DioSn-**2**), and 2 μM (TriSn-**3**, TriSn-**4**), in triplicate across three independent experiments. MRC-5 cells (1 × 10⁵ cells/mL) were treated under the same conditions, with initial highest concentrations of 20 μM (DioSn-**2**) and 5 μM (TriSn-**3**, TriSn-**4**), while DioSn-**1** was excluded due to its inactivity in A549 cells. Cisplatin (50 μM) served as the positive control, and untreated cells as the negative control. After 24 hours, 20 μL of MTT was added and incubated for 4 hours at 37°C with 5% CO₂. The supernatant (180 μL) was removed, followed by the addition of an equal volume of DMSO. Absorbance was measured at 570 nm using a microplate reader. Data were graphically analyzed to determine IC₅₀ values.

### Cell Morphological Changes

Cell morphology assessment followed [16]. A549 cells (5 × 10⁴ cells/mL) were cultured in 6-well plates and treated for 24 hours at IC₅₀ concentrations: 1.86 μM (DioSn-**2**), 0.52 μM (TriSn-**3**), 0.57 μM (TriSn-**4**), and 32 μM (cisplatin). Untreated cells served as the negative control. Morphological changes were observed under an inverted microscope.

### Apoptotic Detection via Annexin V-FITC/PI Staining

The mode of cell death was assessed following Chan et al. [23]. A549 cells (5 × 10⁴ cells/mL) were treated with DioSn-**2**, TriSn-**3**, TriSn-**4**, and cisplatin for 24 hours. Cells were harvested by centrifugation (1200 rpm, 5 min), washed with chilled PBS, and resuspended in 100 μL Annexin binding buffer. Annexin V-FITC (2.5 μL) was added, followed by incubation in the dark at room temperature for 12 minutes. PI (5 μL) was then added and incubated for 3 minutes. After adding 300 μL Annexin binding buffer, samples were analyzed using a FACSCanto II Flow Cytometer (BD Bioscience).

### Comet Assay for DNA Damage Detection

The alkaline comet assay, adapted from Singh et al. [24] was used to assess DioSn-**2**-induced DNA damage in A549 cells. Cells were treated at IC₅₀ for 0.5–4 hours, then centrifuged at 1200 rpm for 5 minutes at 4°C. The supernatant was discarded, and cells were washed twice with cold PBS before resuspension in 80 μL of low-melting agarose (Sigma-Aldrich, USA). The mixture was spread onto pre-chilled microscope slides coated with normal-melting agarose and overlaid with a glass slide. After 15 minutes on ice, slides were immersed in cold lysis buffer with Triton-X (Sigma-Aldrich, USA) overnight. Slides were then placed in an alkaline buffer (pH >13) for 20 minutes before electrophoresis at 25 V and 300 mA for 20 minutes. After three washes with neutralizing buffer (pH 7.5), slides were stained with 50 μL of ethidium bromide (50 μg/mL, Fisher Scientific, Britain). Stained slides were stored in a dark, buffer-dampened container for up to a month. DNA damage was assessed using a fluorescence microscope (Olympus BX51TR, USA) and analyzed via CometScore software (Version 28.0, Tritek Inc.), focusing on the DNA tail moment parameter.

### ROS and Mitochondrial Membrane Potential Assessment via DHE and TMRE

Intracellular ROS levels and mitochondrial membrane potential (Δψm) were assessed using dihydroethidium (DHE) and tetramethylrhodamine ethyl ester (TMRE) staining. A549 cells were cultured in T-25 flasks and treated with DioSn-**2** at its IC₅₀ concentration for 0.5, 1, 2, and 4 hours. The experiment lasted two days: Day 1 for cell seeding and Day 2 for treatment and staining. After overnight incubation, cells were treated with DioSn-**2** in a staggered manner, beginning with the longest incubation time (4 > 2 > 1 > 0.5 hours), alongside cisplatin (positive control) or left untreated (negative control). At the end of the longest incubation period, floating cells and media from each flask were collected into 15 mL centrifuge tubes. Flasks were rinsed twice with PBS, and the rinses were pooled with the collected media. Trypsin-EDTA (1 mL) was added to detach adherent cells, followed by neutralization with complete media. The cell suspension was centrifuged at 1200 rpm for 5 minutes at 4°C to obtain a pellet. The pellet was resuspended in 1 mL of serum-free media and transferred to a microcentrifuge tube. Dihydroethidium (DHE, 10 mM, 1 μL) or Tetramethylrhodamine ethyl ester (TMRE, 50 μM, 1 μL) was added, and samples were incubated in the dark at 37°C for 15 minutes. After incubation, cells were centrifuged at 1200 rpm for 5 minutes at 4°C, washed with cold PBS, and recentrifuged. The final pellet was resuspended in 600 μL of cold PBS and transferred to Falcon tubes for analysis using a BD FACSCanto II flow cytometer.

### NAC Pre-Treatment and ROS-Associated Cytotoxicity

Intracellular ROS levels are often recognized as a key factor contributing to cytotoxicity and leading to cell death. To determine the role of oxidative stress, an experiment using *N*-acetyl-*L*-cysteine (NAC) was conducted. NAC is known for its ability to neutralize ROS and prevent DNA and protein damage caused by oxidative stress. Therefore, cells were pre-treated with 5 mM NAC for 1 hour before being exposed to DioSn-**2** at its IC₅₀ concentration for 24 hours. Following treatment, cells were harvested and analyzed using the Annexin V-FITC/PI assay to assess the involvement of ROS in cytotoxicity.

### Apoptotic Detection via Caspase-8, -9, and -3 Activation

Caspase activation was assessed using the in situ CaspaTag™ assay kit for caspase-8, caspase-9, and caspase-3 to identify the apoptotic pathway triggered by DioSn-**2** in A549 human lung carcinoma cells. A549 cells (5 × 10⁴ cells/mL) were treated with DioSn-**2** and cisplatin (positive control) for 4 and 24 hours, while untreated cells served as a negative control. Following incubation, floating cells and media were collected, and adherent cells were detached using trypsin-EDTA, neutralized with complete media, and pooled with the collected media. After centrifugation at 1200 rpm for 5 minutes, the cell pellet was resuspended in 10 μL of 30X FLICA reagent (specific for caspase-8, -9, and -3) and incubated at 37°C for 1 hour in a 5% CO₂ incubator. The cells were then washed twice with 1X washing buffer, resuspended in 400 μL of 1X washing buffer, and analyzed using a BD FACSCanto II flow cytometer.

### Statistical Analysis

Statistical analyses were conducted using IBM SPSS version 28.0. One-way ANOVA was performed to compare the percentage of viable A549 cells across different compound concentrations and to assess viable, apoptotic, and necrotic cell populations at IC₅₀ concentrations. Data were presented as mean ± standard error of the mean (S.E.M.) from three independent experiments, with p-values < 0.05 considered statistically significant.

## RESULTS

### Synthesize and characterization of compounds

The *in-situ* method is an efficient approach for producing dithiocarbamic acid, as solid synthesis becomes challenging above 4°C. This exothermic reaction between carbon disulfide and secondary amines may produce byproducts at high temperatures [25]. Nevertheless, the method consistently achieves high success rates, with yields exceeding 70%. Table 1 presents the physical data for DioSn and TriSn compounds.

**Table 1.** Physical and Elemental Data of DioSn and TriSn compounds.

| Molecular formula | Colour | Yield | Melting Point (°C) | Experimental analysis found |  |  |  |
| --- | --- | --- | --- | --- | --- | --- | --- |
|  |  |  |  | (Calc.)<br>% C | % H | % N | % S |
| $(\text{CH}_3)_2\text{Sn}[\text{S}_2\text{CN}(\text{CH}_3(\text{C}_6\text{H}_5)(\text{CH}_2))_2]$ | White | 80% | 141.9–<br>144.5 | **44.37**<br>43.40 | **4.84**<br>4.79 | **5.17**<br>5.03 | **23.69**<br>23.80 |
| $(\text{C}_6\text{H}_5)_2\text{Sn}[\text{S}_2\text{CN}(\text{CH}_3(\text{C}_6\text{H}_5)(\text{CH}_2))_2]$ | White | 89% | 166.0–<br>168.0 | **54.14**<br>53.58 | **4.54**<br>4.56 | **4.21**<br>3.15 | **19.27**<br>16.17 |
| $(\text{C}_6\text{H}_5)_3\text{Sn}[\text{S}_2\text{CN}(\text{CH}_3(\text{C}_6\text{H}_5)(\text{CH}_2))]$ | White | 89% | 82.8–<br>84.5 | **59.36%**<br>59.24% | **4.61%**<br>4.77% | **2.56%**<br>1.49% | **11.74**<br>9.01% |
| $(\text{C}_6\text{H}_5)_3\text{Sn}[\text{S}_2\text{CN}(\text{CH}_3\text{CH}_2(\text{C}_6\text{H}_5)(\text{CH}_2))]$ | White | 70% | 92–94.8 | **60.02%**<br>60.39% | **4.86%**<br>4.94% | **2.50%**<br>1.45% | **11.44**<br>10.53% |

### FTIR absorbance peaks for DioSn and TriSn compounds

Key vibrational bands in dithiocarbamate complexes include υ(C**<u>••••</u>**N), υ(C**<u>••••</u>**S), υ(Sn–C), and υ(Sn–S). The thioureide band υ(C**<u>••••</u>**N) (1479–1497 cm⁻¹) and υ(C**<u>••••</u>**S) (950–998 cm⁻¹) indicate electron delocalization in the -NCS₂ group [17,26]. Additionally, absorption bands at 445–569 cm⁻¹ correspond to Sn-C vibrations, while those at 369–388 cm⁻¹ are attributed to Sn-S vibrations. A summary of the key FTIR absorbance peaks for DioSn and TriSn compounds is provided in Table 2.

**Table 2.** FTIR absorbance peaks for DioSn and TriSn compounds.

| Compounds | Wavelength ($\text{cm}^{-1}$) | | | | | |
| --- | --- | --- | --- | --- | --- | --- |
| | $\nu(\text{C}\cdots\text{N})$ | $\nu(\text{C}\cdots\text{S})$ | $\nu(\text{Sn}-\text{C})$ | $\nu(\text{Sn}-\text{S})$ | $\nu(\text{N}-\text{C})$ | $\nu(\text{C}-\text{H})$ |
| **DioSn-1** | 1488 | 998 | 553 | 374 | 1269 | 2911 |
| **DioSn-2** | 1494 | 950 | 569 | 383 | 1256 | 3064 |
| **TriSn-3** | 1497 | 979 | 496 | 388 | 1254 | 3056 |
| **TriSn-4** | 1479 | 997 | 445 | 369 | 1077 | 3044 |

### ¹³C and ¹¹⁹Sn NMR data for DioSn and TriSn compounds

¹³C NMR analysis identified the thioureide carbon resonance at 197–201 ppm and the methylene carbon at 58.42–61.64 ppm. The methyl carbon attached to the *N*-methyl-*N*-benzyl dithiocarbamate ligand exhibited chemical shifts ranging from δ 41.77 to δ 42.90 ppm. In contrast, the methyl carbon of the ethyl group in the *N*-ethyl-*N*-benzyl dithiocarbamate ligand resonated at a lower frequency (downfield) at δ 11.84 ppm. Meanwhile, carbon signals observed between δ 127 and δ 151 ppm correspond to the aromatic phenyl group. The geometric structures of the compounds were confirmed by ¹¹⁹Sn NMR, revealing their dependence on the ligand type and surrounding environment, consistent with previous studies [14]. Specifically, DioSn-**1** and DioSn-**2** exhibited octahedral and pentagonal bipyramidal geometries, respectively, while TriSn-**3** and TriSn-**4** displayed a trigonal bipyramidal geometry. Table 3 summarizes the ¹³C and ¹¹⁹Sn NMR data for the DioSn and TriSn compounds.

**Table 3.**
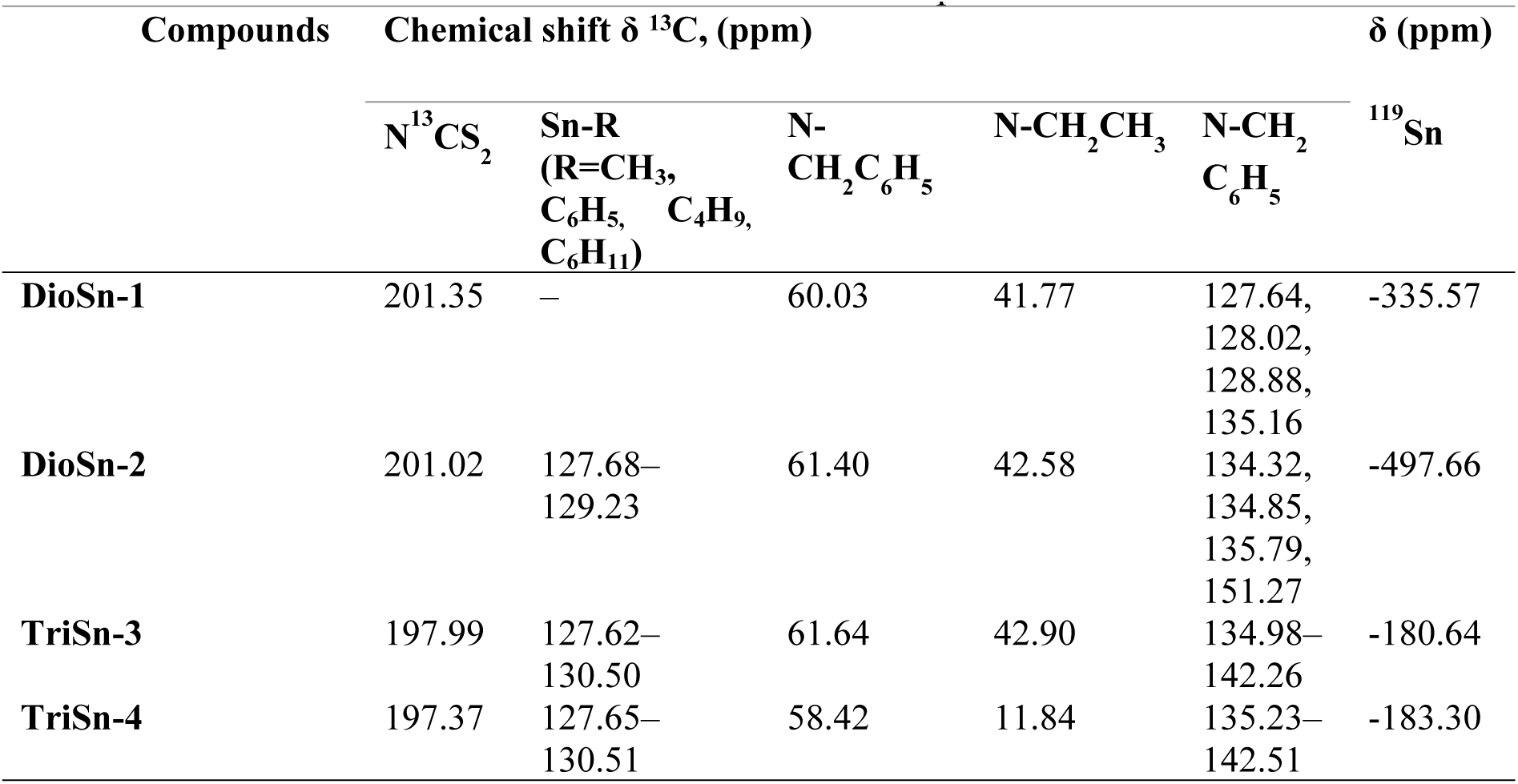
¹³C and ¹¹⁹Sn NMR data for DioSn and TriSn compounds.

| Compounds | Chemical shift $\delta$ $^{13}\text{C}$, (ppm) | | | | | $^{119}\text{Sn}$<br>$\delta$ (ppm) |
| --- | --- | --- | --- | --- | --- | --- |
| | $\text{N}^{13}\text{CS}_2$ | Sn-R<br>(R=CH<sub>3</sub>,<br>C<sub>6</sub>H<sub>5</sub>, C<sub>4</sub>H<sub>9</sub>,<br>C<sub>6</sub>H<sub>11</sub>) | N-CH<sub>2</sub>C<sub>6</sub>H<sub>5</sub> | N-CH<sub>2</sub>CH<sub>3</sub> | N-CH<sub>2</sub><br>C<sub>6</sub>H<sub>5</sub> | |
| DioSn-1 | 201.35 | – | 60.03 | 41.77 | 127.64,<br>128.02,<br>128.88,<br>135.16 | -335.57 |
| DioSn-2 | 201.02 | 127.68–<br>129.23 | 61.40 | 42.58 | 134.32,<br>134.85,<br>135.79,<br>151.27 | -497.66 |
| TriSn-3 | 197.99 | 127.62–<br>130.50 | 61.64 | 42.90 | 134.98–<br>142.26 | -180.64 |
| TriSn-4 | 197.37 | 127.65–<br>130.51 | 58.42 | 11.84 | 135.23–<br>142.51 | -183.30 |

### Recrystallization, Molecular structure, and Crystal Packing Analyses

Three compounds (DioSn-**1**, TriSn-**3**, and TriSn-**4**) were successfully recrystallized, yielding single crystals. The crystallographic data for DioSn-**1** and TriSn-**3** are reported in this study, while those for TriSn-**4** were previously published [21]. DioSn-**1** crystals measured 0.29 × 0.07 × 0.05 mm, while TriSn-**3** had a slightly smaller size of 0.18 × 0.09 × 0.04 mm. Their molecular weights were 541.36 g and 546.29 g, respectively. Both crystallized in the monoclinic system (P2₁/n). Unit cell parameters for DioSn-**1** were *a* = 12.0888(2) Å, *b* = 9.0576(1) Å, *c* = 20.9040(2) Å, *β* = 92.810(1)°, while for DioSn-**2**, they were *a* = 6.7703(1) Å, *b* = 12.0476(1) Å, *c* = 29.5565(1) Å, *β* = 96.361(1)°. X-ray diffraction was performed using a Rigaku XtaLAB Synergy Dualflex AtlasS diffractometer with CuKα radiation (λ = 1.541 Å). Data collection, cell refinement, and reduction were conducted using CrysAlis PRO [27]. Molecular structures were determined via SHELXTL [28] using direct methods and refined through full-matrix least-squares on F² with anisotropic displacement. Empirical absorption correction was applied using SADABS [27], while geometric calculations were performed using PLATON [29]. Non-hydrogen atoms were refined anisotropically, while hydrogen atoms were placed geometrically (isotropic displacement) and refined via the riding model. Methyl groups were treated using a rotating model. Table 4 summarizes the key refinement parameters, and crystallographic data have been deposited in the Cambridge Structural Database (CSD). Meanwhile, table 5, 6 and 7 list the selected bond lengths and angles for the molecular structures of DioSn-**1**, TriSn-**3**, and TriSn-**4**, respectively. Ortep plots and supramolecular arrangements of DioSn-**1**, TriSn-**3**, and TriSn-**4** are shown in Figures 1–3.

**Figure 1.**
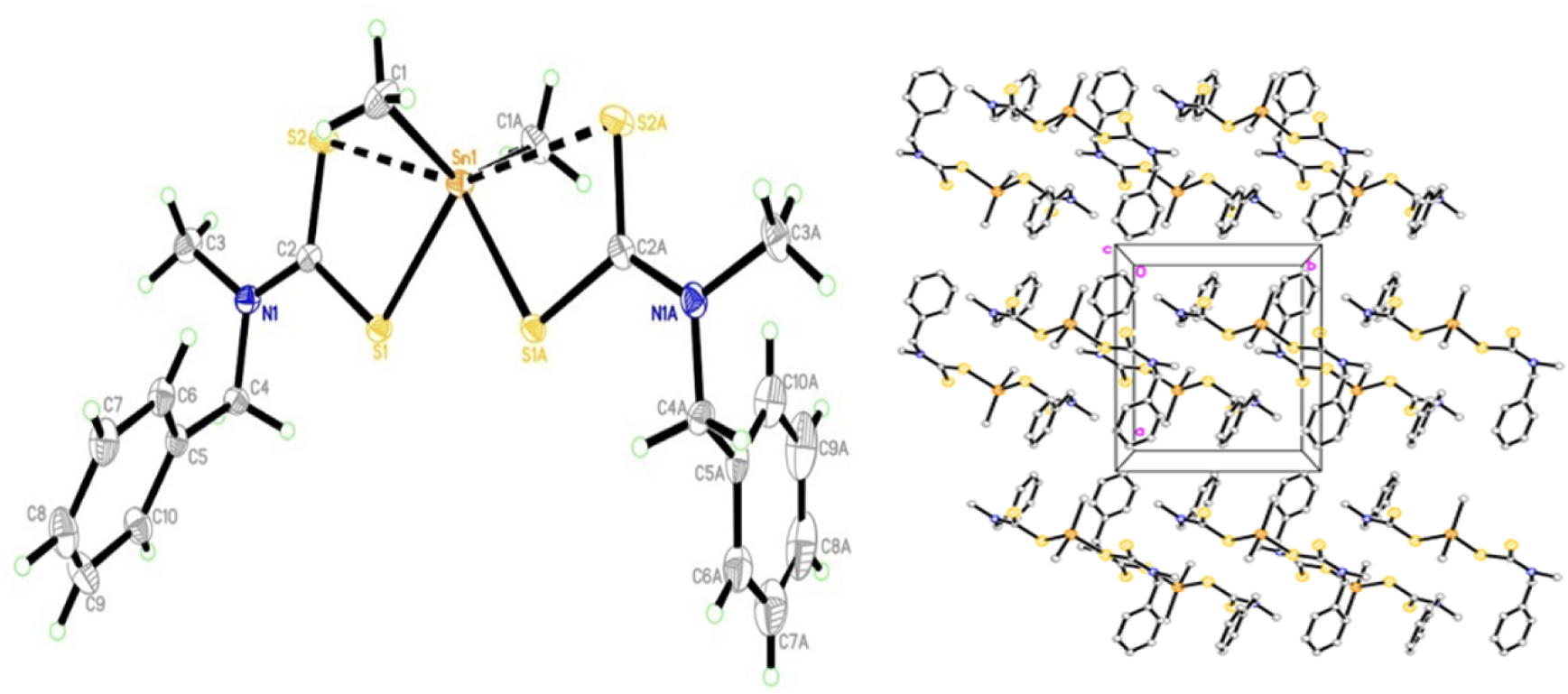
ORTEP plot of DioSn-**1** at 50% ellipsoid probability with atomic numbering. Supramolecular layers are indicated by grey dashed lines.

**Figure 2.**
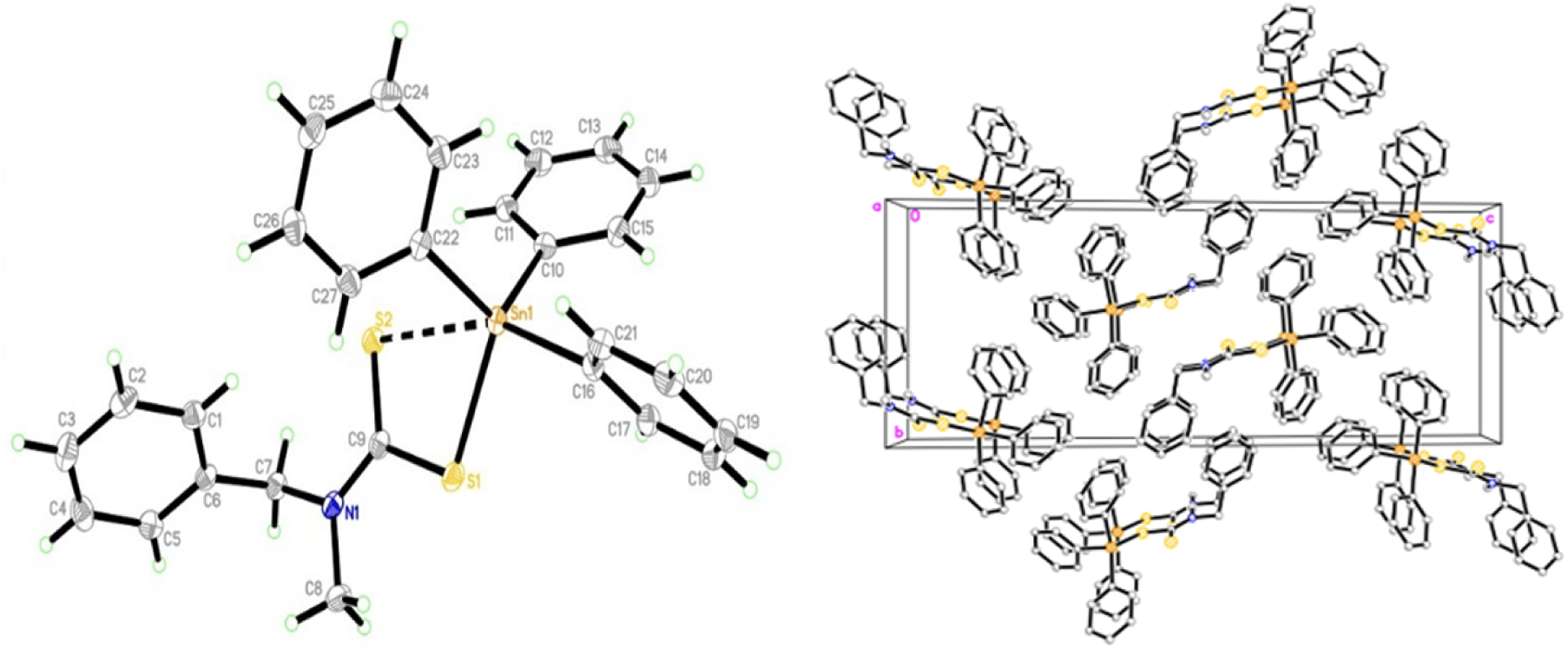
ORTEP plot of TriSn-**3** at 50% ellipsoid probability with atomic numbering. Supramolecular layers linked by C-H···π interactions are shown as grey dashed lines.

**Figure 3.**
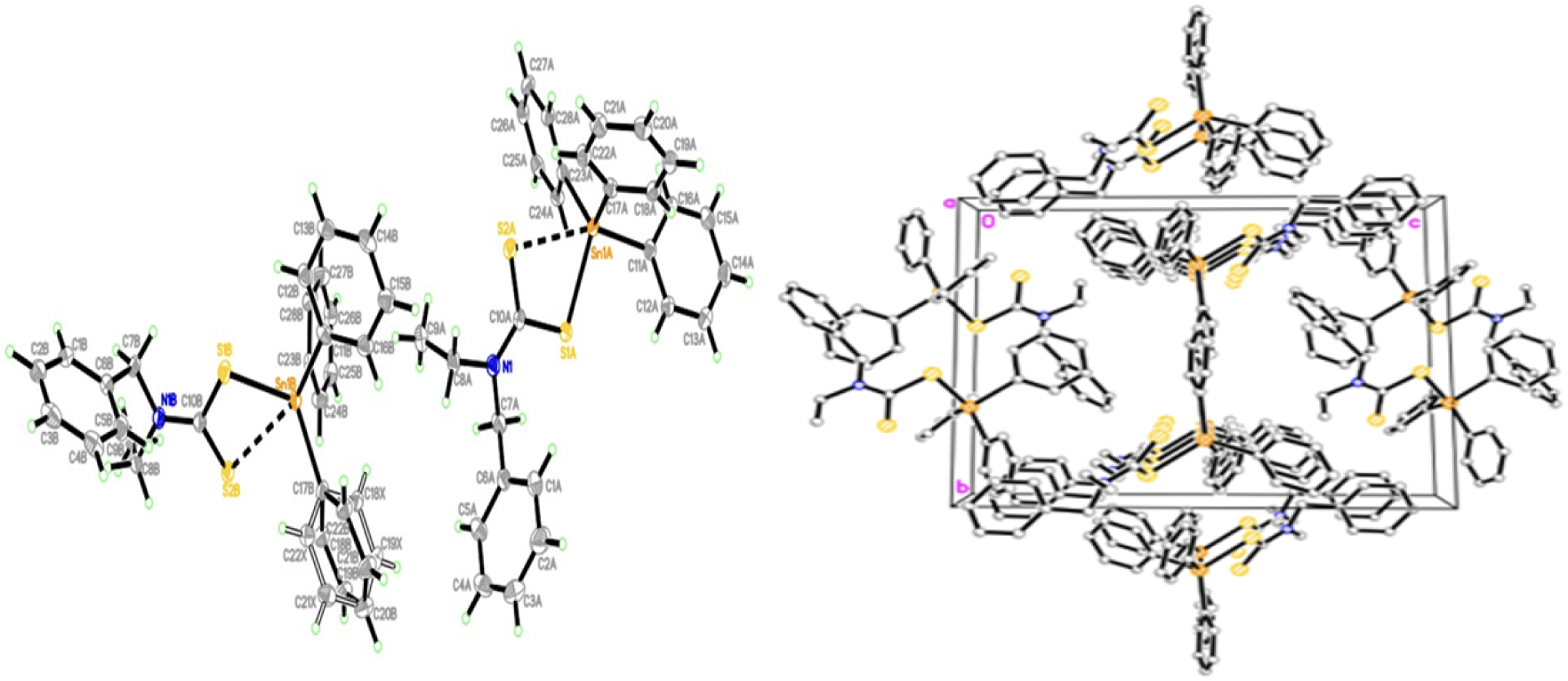
ORTEP plot of TriSn-**4** at 50% ellipsoid probability with atomic numbering. Supramolecular layers linked by C-H···π interactions are shown as grey dashed lines [21].

**Table 4.**
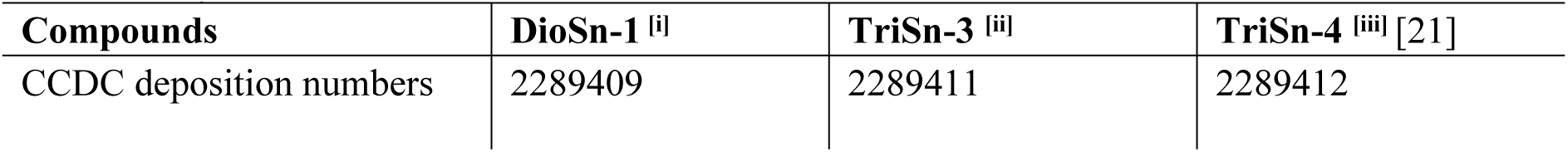
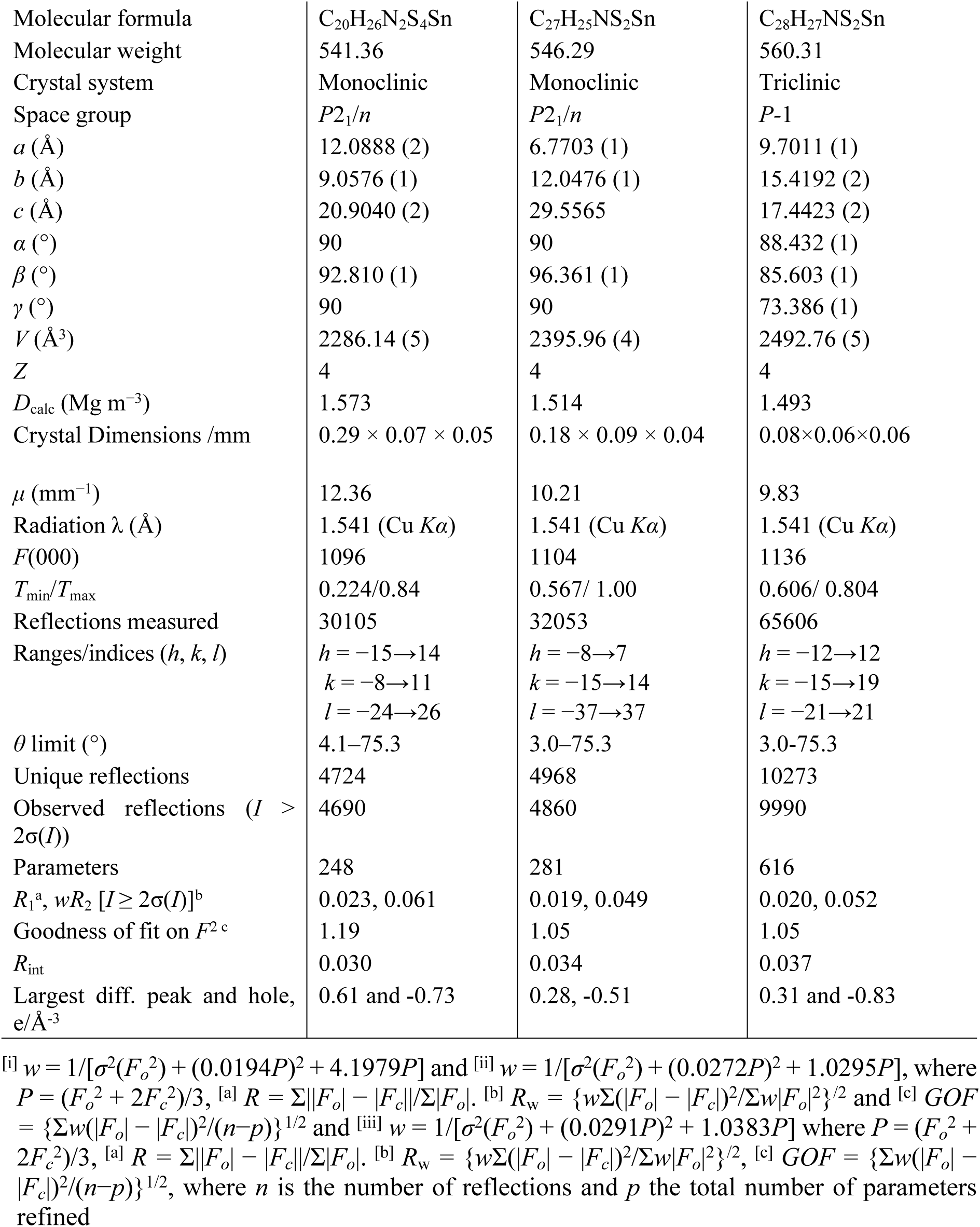
Crystal data and structure refinement of DioSn-1, TriSn-3 and TriSn-4.

**Table 5:**
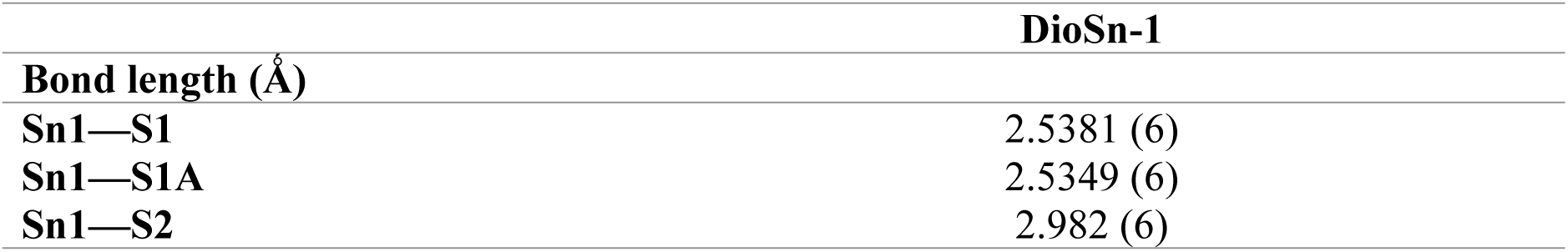
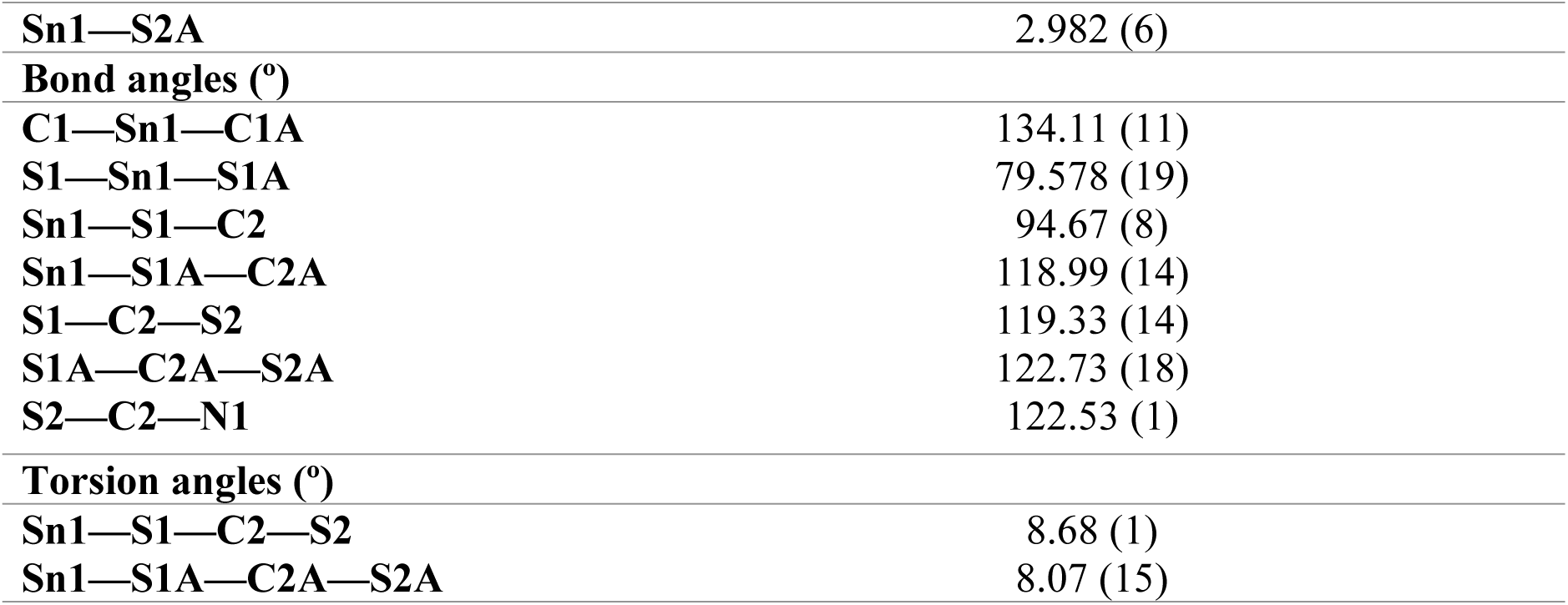
The selected geometric parameters of DioSn-1.

**Table 6:** The selected geometric parameters of TriSn-3.

|  | **TriSn-3** |
| --- | --- |
| **Bond length (Å)** |  |
| **Sn1—S1** | 2.4724 (4) |
| **Sn1—S2** | 3.020 (4) |
| **Bond angles (°)** |  |
| **C9—S1—Sn1** | 96.06 (5) |
| **S2—C9—S1** | 118.85 (9) |
| **N1—C9—S1** | 117.53 (1) |
| **Torsion angles (°)** |  |
| **Sn1—S1—C9—S2** | 11.23 (9) |

### Cytotoxicity of DioSn and TriSn compounds in A549 and MRC-5 cells

The MTT assay was performed to determine the IC₅₀ values of DioSn and TriSn compounds in A549 cells after 24 hours of treatment (Figures 4–6). Tables 9 summarize these IC₅₀ values. DioSn-**2**, TriSn-**3**, and TriSn-**4** reduced cell viability at concentrations as low as 5 μM and 2 μM, respectively, while DioSn-**1** remained inactive even at 50 μM. The IC₅₀ values were 1.86 μM for DioSn-**2**, 0.57 μM for TriSn-**3**, and 0.52 μM for TriSn-**4**. As a positive control, cisplatin had an IC₅₀ of 32 μM after 24 hours (Figure 7). Table 2 presents the IC₅₀ values and selectivity indices (SI) of all tested compounds, highlighting DioSn-**2** as the most selectively cytotoxic toward A549 cells.

**Figure 4.**
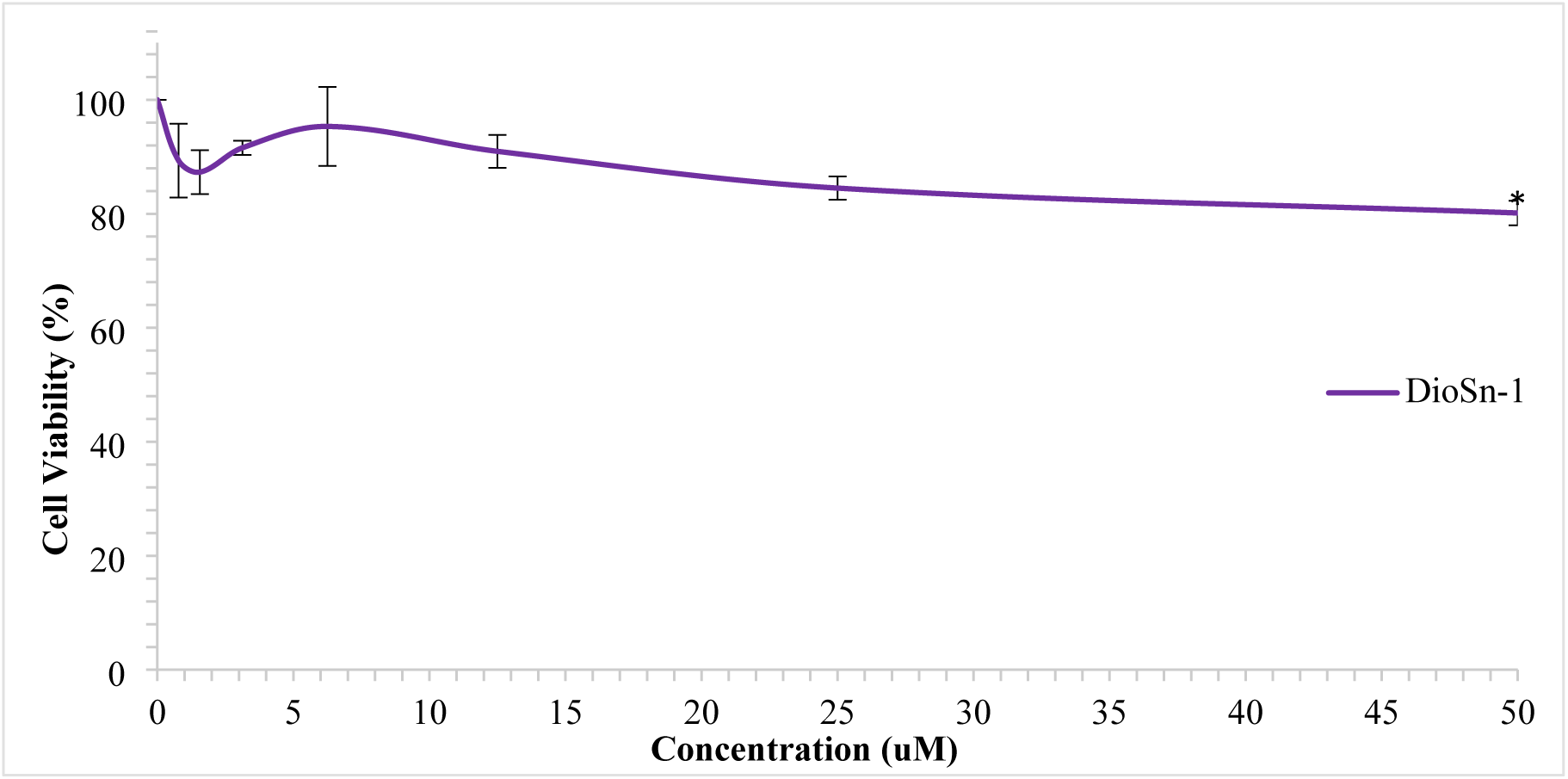
The IC_50_ value on A549 cells after treated with DioSn-1 for 24 hours of treatment. Data are presented as mean ± S.E.M from three independent experiments. *Significant differences at the *p* < 0.05 compared to negative control

**Figure 5.**
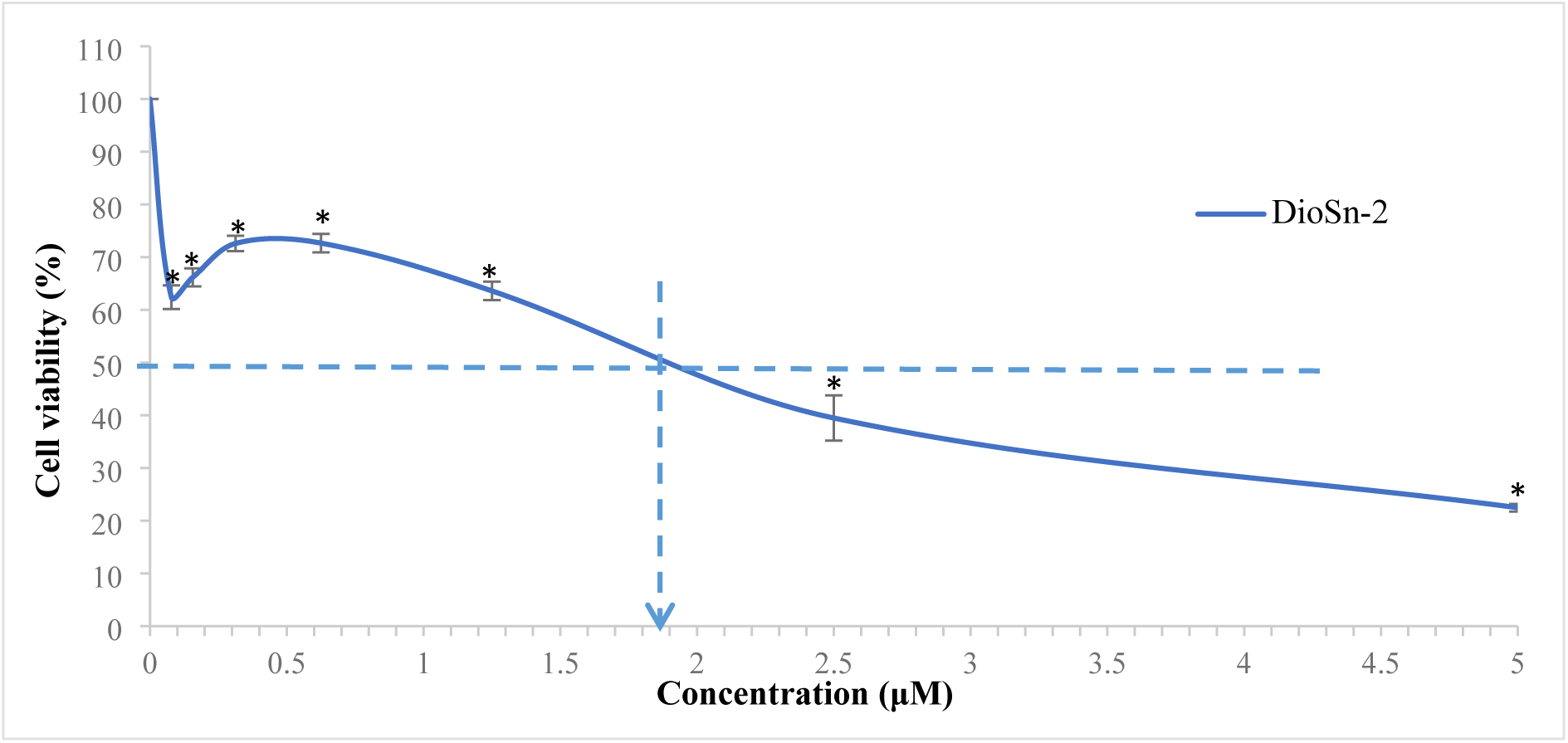
The IC_50_ value on A549 cells after treated with DioSn-2 for 24 hours of treatment. Data are presented as mean ± S.E.M from three independent experiments. *Significant differences at the *p* < 0.05 compared to negative control

**Figure 6.**
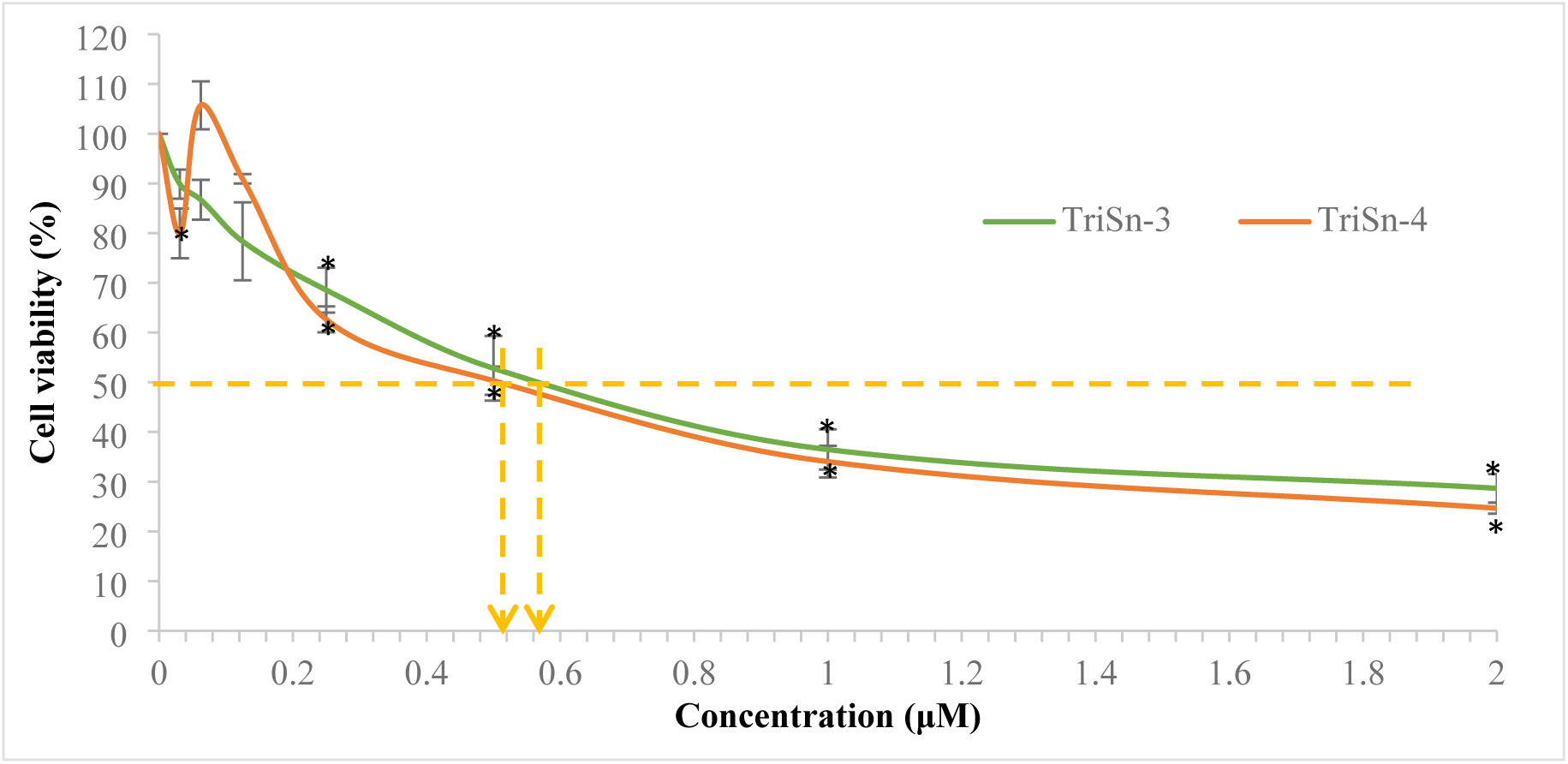
The IC_50_ value on A549 cells after treated with TriSn-3 and TriSn-4 for 24 hours of treatment. Data are presented as mean ± S.E.M from three independent experiments. *Significant differences at the *p* < 0.05 compared to negative control

**Figure 7.**
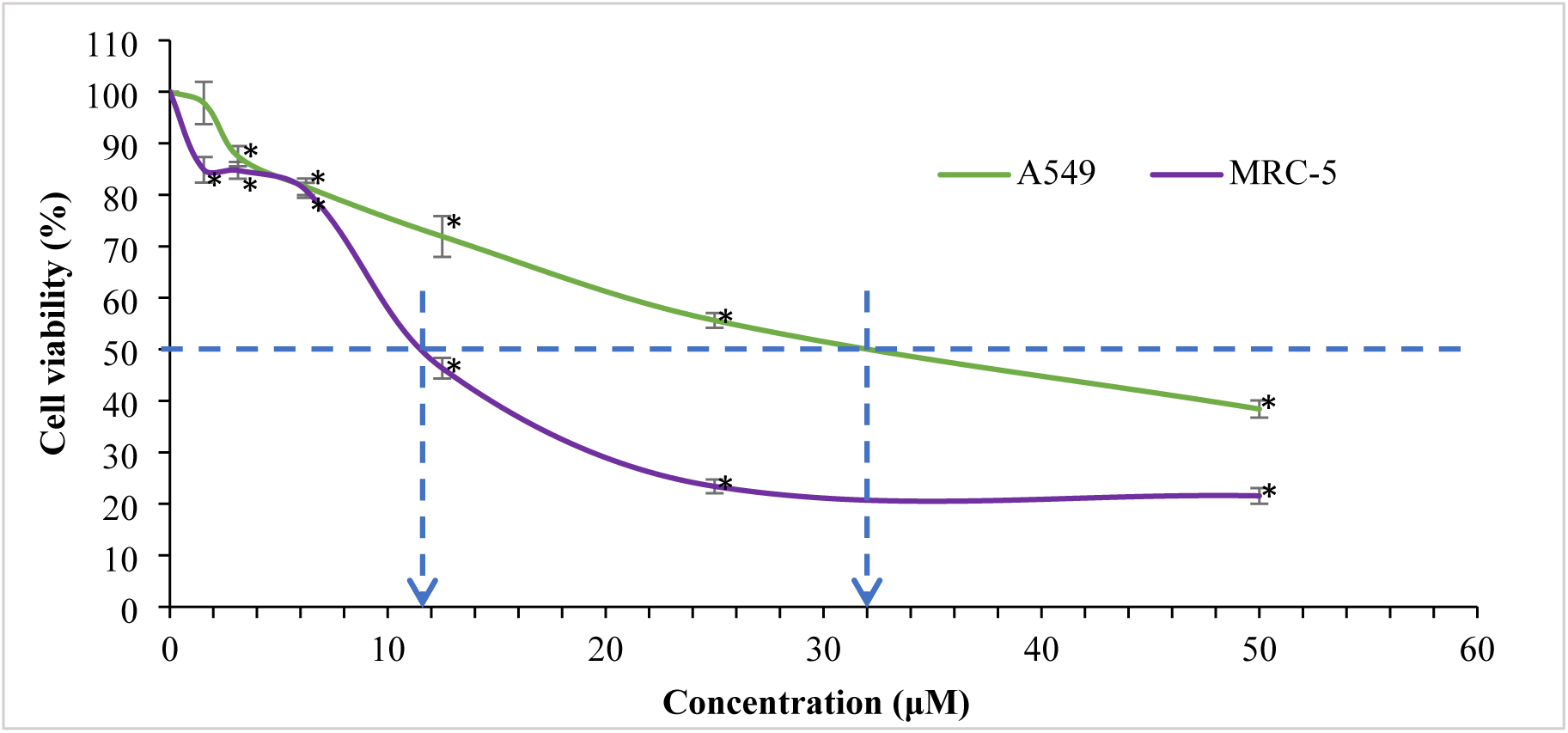
IC₅₀ values of cisplatin in A549 and MRC-5 cells after 24 hours of treatment. Data are presented as mean ± S.E.M from three independent experiments. *Significant differences at the *p* < 0.05 compared to negative control

### Morphological changes of DioSn-2, TriSn-3, and TriSn-4 in A549 cells

Morphological assessments were performed on A549 cells treated with the compounds at their IC_50_ concentrations from the MTT assay for 24 hours, including cisplatin. Observations under an inverted microscope (Figure 8) revealed characteristic apoptotic changes, such as cell shrinkage (S), plasma membrane blebbing (B), and apoptotic body formation (AB). Apoptosis, a programmed cell death process, is marked by nuclear chromatin condensation, cell shrinkage, membrane blebbing, and apoptotic body formation [30].

**Figure 8.**
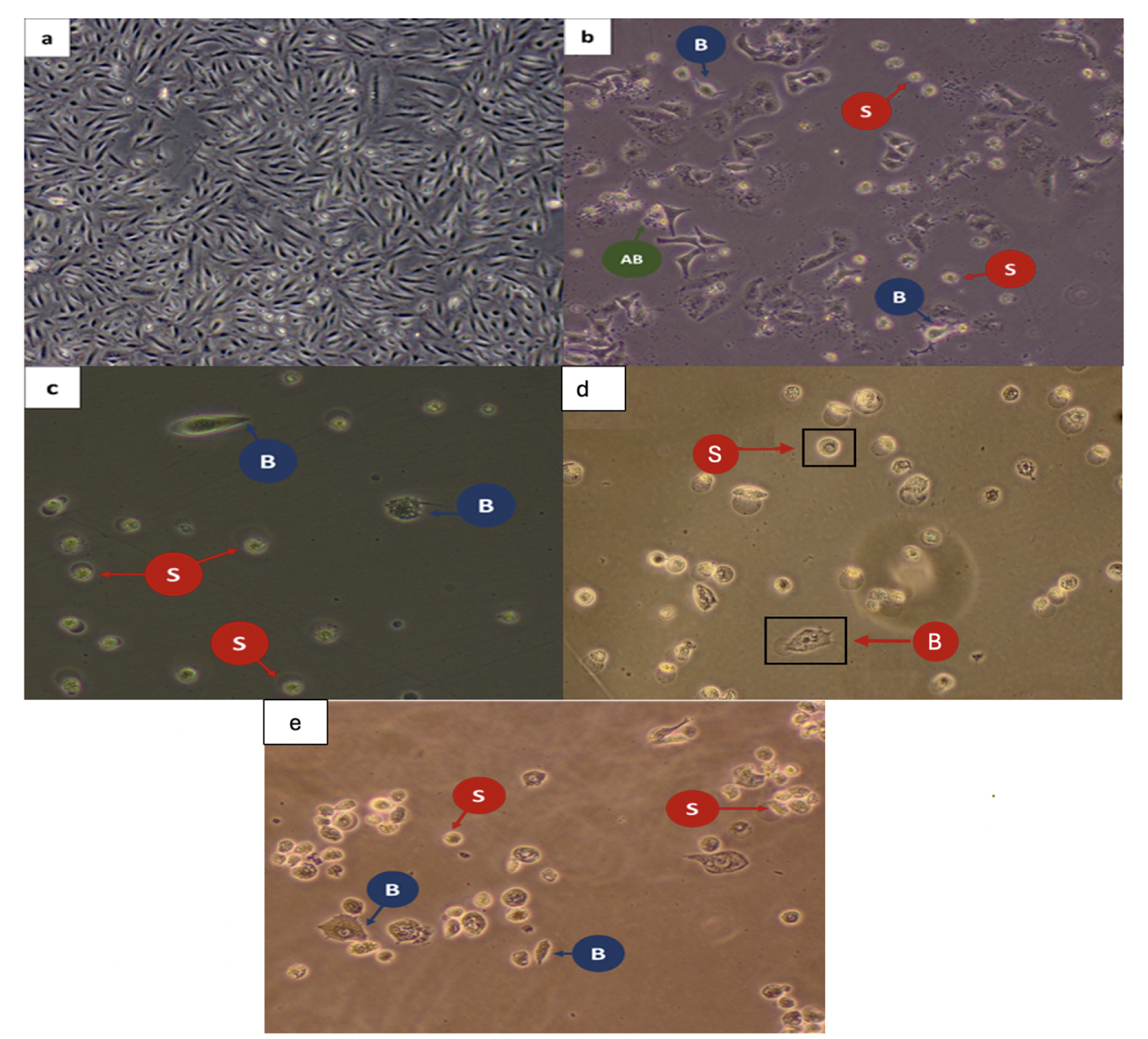
Morphological changes in A549 cells (40×): (a) Untreated control, (b) Cisplatin-treated (24 h), (c) DioSn-2 treated (24 h), (d) TriSn-3 treated (24 h), and (e) TriSn-4 treated (24. **h)**

### DioSn-2, TriSn-3 and TriSn-4 compounds induced apoptosis in A549 cells

Annexin V-FITC/PI staining was used to assess apoptosis and necrosis, validating the cell death mechanism induced by the compounds in A549 cells [31]. Figure 9 shows apoptotic rates at IC_50_ concentrations after 24 hours: DioSn-**2** (59.90 ± 3.76%), TriSn-**3** (58.27 ± 0.83%), and TriSn-**4** (91.40 ± 0.52%). All compounds significantly induced apoptosis (*p* < 0.05) compared to the negative control (7.10 ± 0.47%).

**Figure 9.**
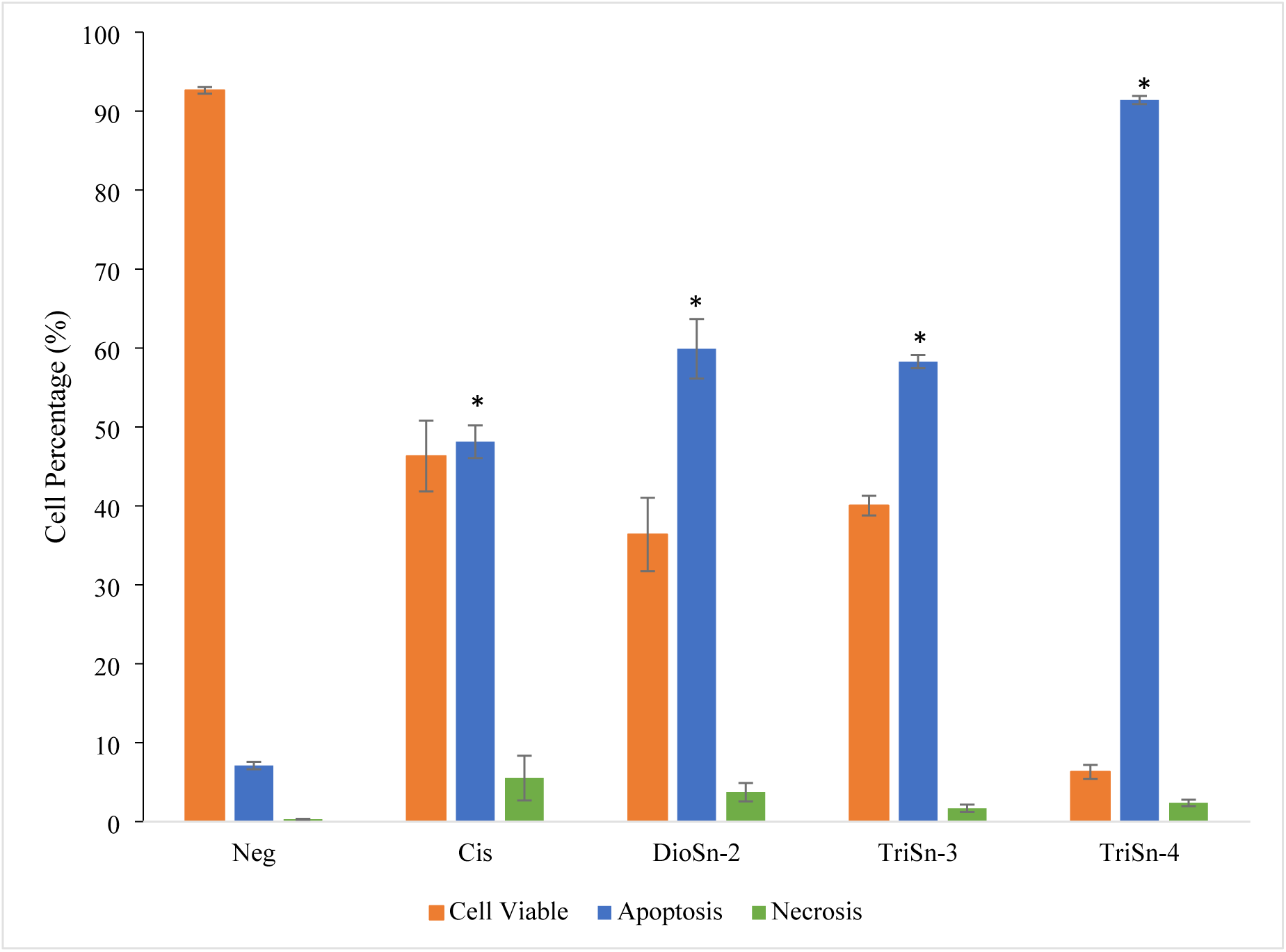
Viable, apoptotic, and necrotic A549 cell percentages after 24-hour treatment at IC_50_ concentrations, assessed via Annexin V-FITC/PI staining. Data are presented as mean ± SEM from three independent experiments. \**p* < 0.05 vs. negative control.

### DNA Damage Induced by DioSn-2 in A549 Cells

DNA damage induced by DioSn-**2** in A549 cells at IC_50_ (1.86 μM) was assessed using the alkaline comet assay. Electrophoresis revealed DNA strand breaks migrating toward the anode, forming a comet-like structure visible under a fluorescence microscope (Figure 10A). Figure 10B shows that DioSn-**2** induced DNA damage at the IC_50_ concentration as early as 30 minutes, increasing with exposure time. This was evident from nuclear images, where longer treatment durations resulted in extended comet tails. Furthermore, DNA damage at all four time points showed significantly higher tail moments than the positive control (1.06 ± 0.02) and a notable difference (*p* < 0.05) compared to the negative control (0.52 ± 0.03). The highest tail moment was observed at 4 hours (2.18 ± 0.10), followed by 2 hours (1.48 ± 0.21), 1 hour (1.38 ± 0.04), and 30 minutes (1.26 ± 0.07).

**Figure 10.**
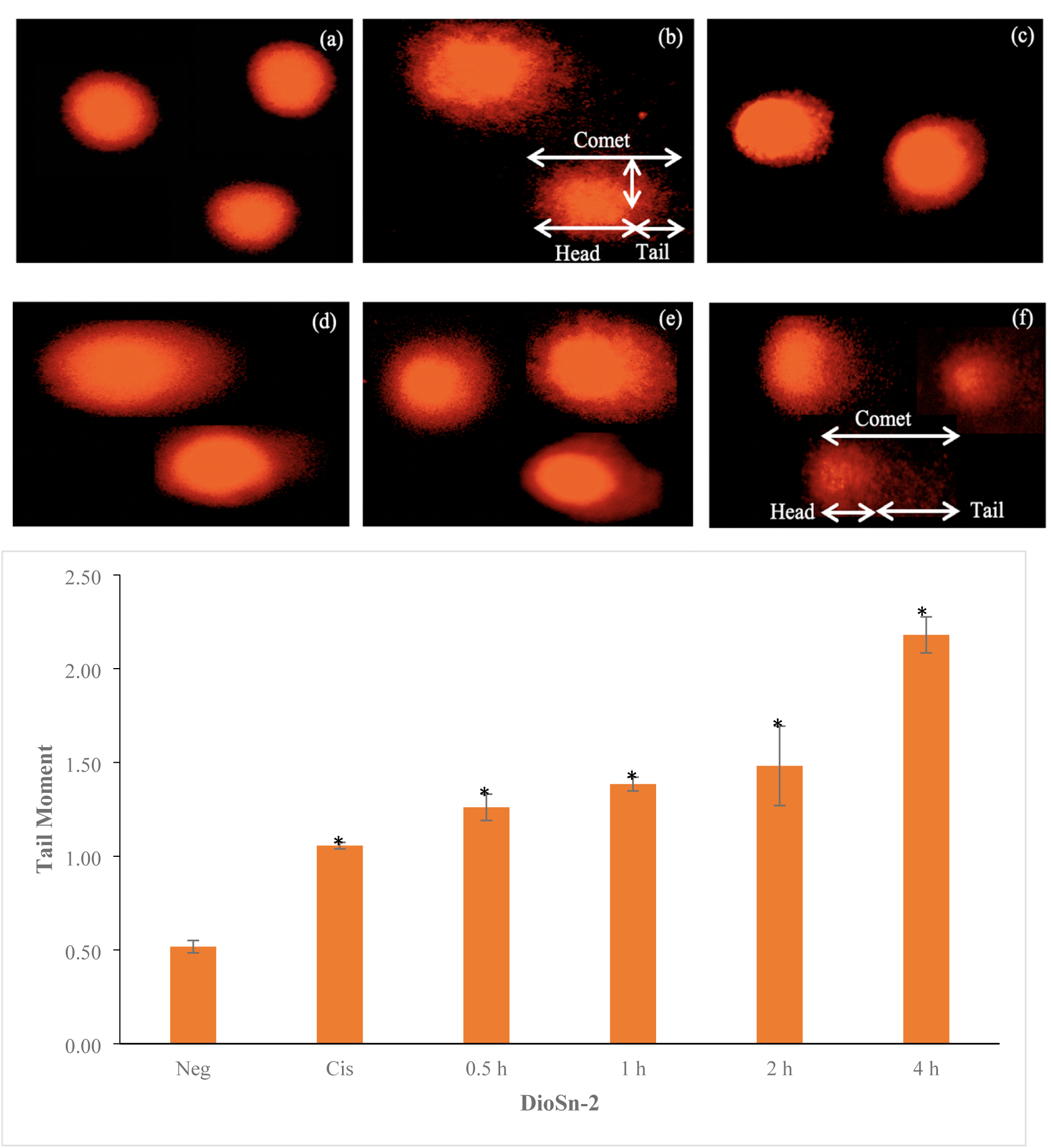
(A) Nuclear images of A549 cells: (a) Negative control (untreated), (b) Cisplatin-treated, (c) DioSn-2 treatment for 0.5 h, (d) 1 h, (e) 2 h, and (f) 4 h. (B) DNA damage induction in A549 cells by DioSn-2 at IC_50_ (1.86 μM). Data are presented as mean ± SEM from three independent experiments. \**p* < 0.05 vs. negative control.

### Δψm Loss Induced by DioSn-2 in A549 Cells

DioSn-**2**-induced mitochondrial membrane potential (Δψm) depolarization in A549 cells was assessed using the TMRE staining assay, as mitochondria regulate pro-apoptotic and anti-apoptotic factor release upon apoptotic stimulation. The bar chart in Figure 11 shows a time-dependent Δψm reduction in cells treated at the IC_50_ concentration. At 30 minutes, a slight but non-significant (*p* > 0.05) increase in TMRE-negative cells was observed compared to the negative control, indicating an early Δψm decrease (11.87 ± 0.89%). This reduction became significant (*p* < 0.05) at 1 hour (14.20 ± 0.51%) and continued to decline at 2 hours (14.83 ± 1.48%) and 4 hours (19.07 ± 1.43%) compared to untreated cells. This Δψm loss may contribute to apoptosis induction by DioSn-**2**.

**Figure 11.**
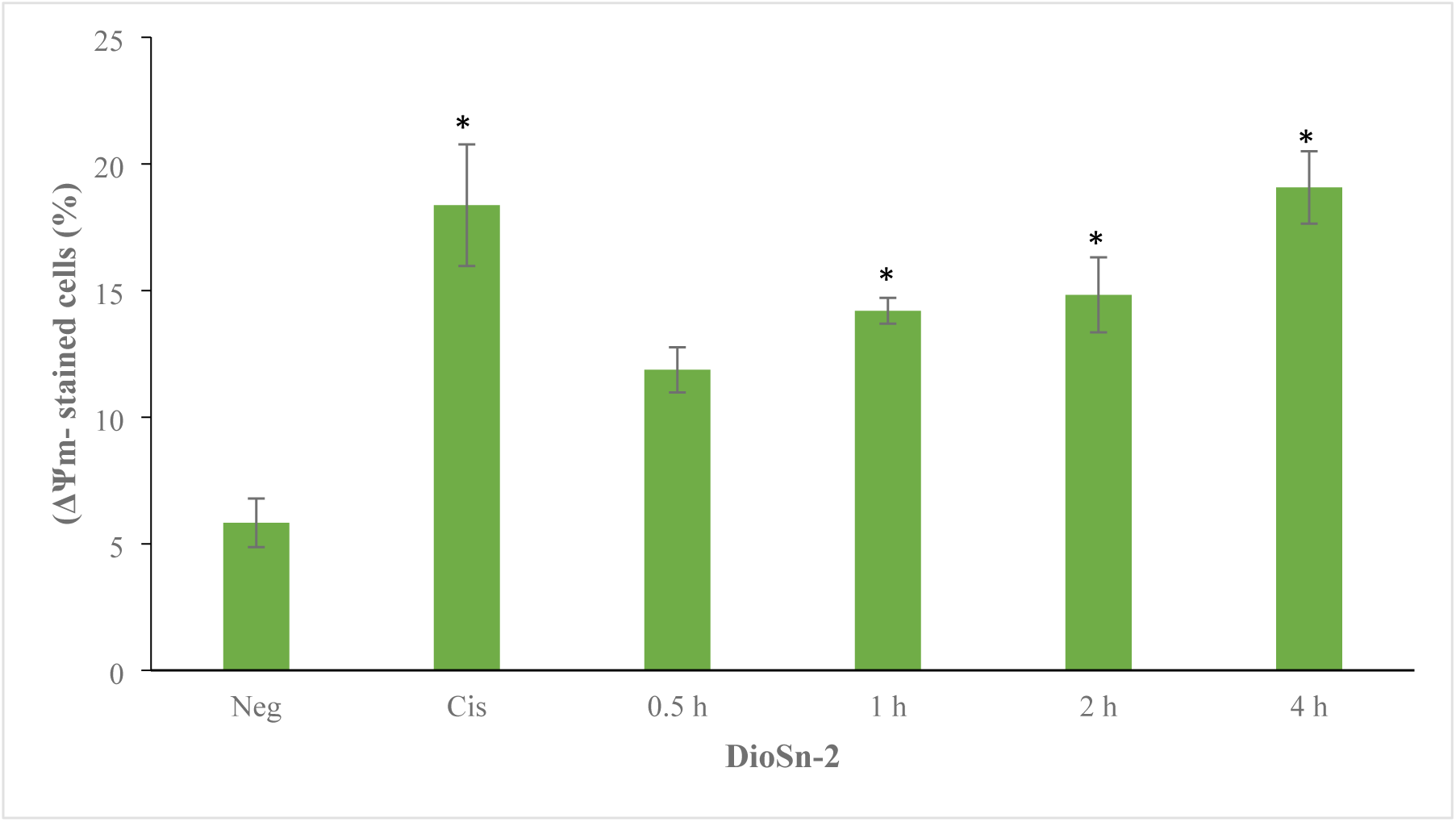
DioSn-2 induces mitochondrial membrane potential (Δψm) depolarization in A549 cells at IC50 (1.86 μM). Data are presented as mean ± SEM from three independent experiments. \**p* < 0.05 vs. negative control.

### ROS-Associated Cytotoxicity of DioSn-2 Confirmed by NAC Pre-Treatment

Loss of Δψm may be linked to disrupted intracellular redox balance, leading to excessive ROS formation. To assess this, intracellular ROS levels were measured using the DHE assay in A549 cells treated with DioSn-**2** at IC_50_ for 30 minutes to 4 hours. As shown in Figure 12A, oxidative stress increased, indicated by a rise in DHE-positive cells as early as 30 minutes, though not significant (*p* > 0.05). ROS levels initially declined after 30 minutes but later increased, indicating DioSn-**2** stimulates superoxide and hydrogen peroxide production, with a significant rise (*p* < 0.05) in DHE-positive cells at 4 hours (29.60 ± 2.72%). To further confirm the role of oxidative stress in DioSn-**2**-induced apoptosis, A549 cells were pre-treated with the synthetic antioxidant *N*-acetyl-*L*-cysteine (NAC, 5 mM) for 1 hour before DioSn-**2** exposure at IC_50_ for 24 hours. Annexin V-FITC/PI staining (Figure 12B) revealed that without NAC, viable cells decreased to 36.37 ± 4.65%, while apoptotic cells significantly increased (*p* < 0.05) to 59.90 ± 3.76%. NAC pre-treatment inhibited apoptosis by 52%, increasing viable cells to 91.13 ± 1.56% and reducing apoptotic cells to 7.43 ± 2.07% (*p* < 0.05). Cisplatin, used as a positive control, also showed significant differences (p < 0.05) between NAC-treated and untreated cells. These findings confirm that oxidative stress from excessive ROS production plays a crucial role in DioSn-**2**-induced apoptosis.

**Figure 12.**
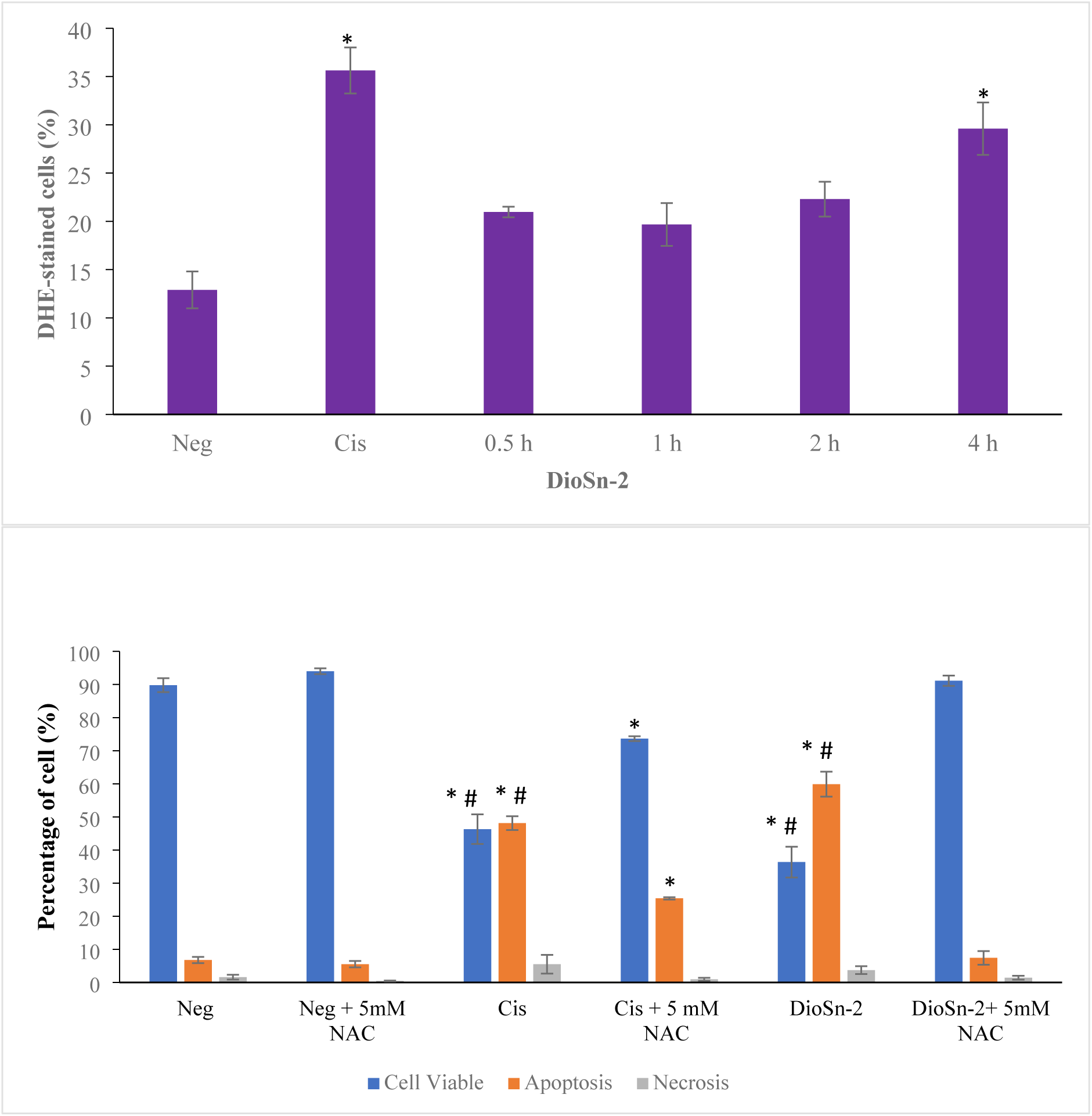
(A) DioSn-2 induces ROS production in A549 cells at IC₅₀ (1.86 μM) (B) Percentage of viable, apoptotic, and necrotic A549 cells after NAC pre-treatment (5 mM). Data represent mean ± S.E.M from three independent experiments. \**p* < 0.05 vs. negative control. #*p* < 0.05 vs. compound + 5 mM NAC.

### Apoptotic Detection via Caspase-8, -9, and -3 Activation in DioSn-2-Treated A549 Cells

Caspase activation assay was performed to determine the apoptotic pathway induced by DioSn-**2** in A549 cells. After 4 hours of treatment, caspase-9 activation (27.73 ± 0.49%) was slightly higher than caspase-8 (26.33 ± 0.24%), both significantly different (*p* < 0.05) from the untreated control. However, after 24 hours, caspase-9 activation increased markedly (44.63 ± 0.86%) compared to caspase-8 (18.67 ± 0.94%), with both being significantly higher (*p* < 0.05) than the negative control (Figure 12). Meanwhile, caspase-3 activation significantly increased after 4 hours (59.20 ± 2.48%) and further increased after 24 hours. Cisplatin, used as a positive control, also showed a significant (*p* < 0.05) increase in caspase-9 and caspase-3 activation after 4 and 24 hours compared to the negative control (Figure 13).

**Figure 13.**
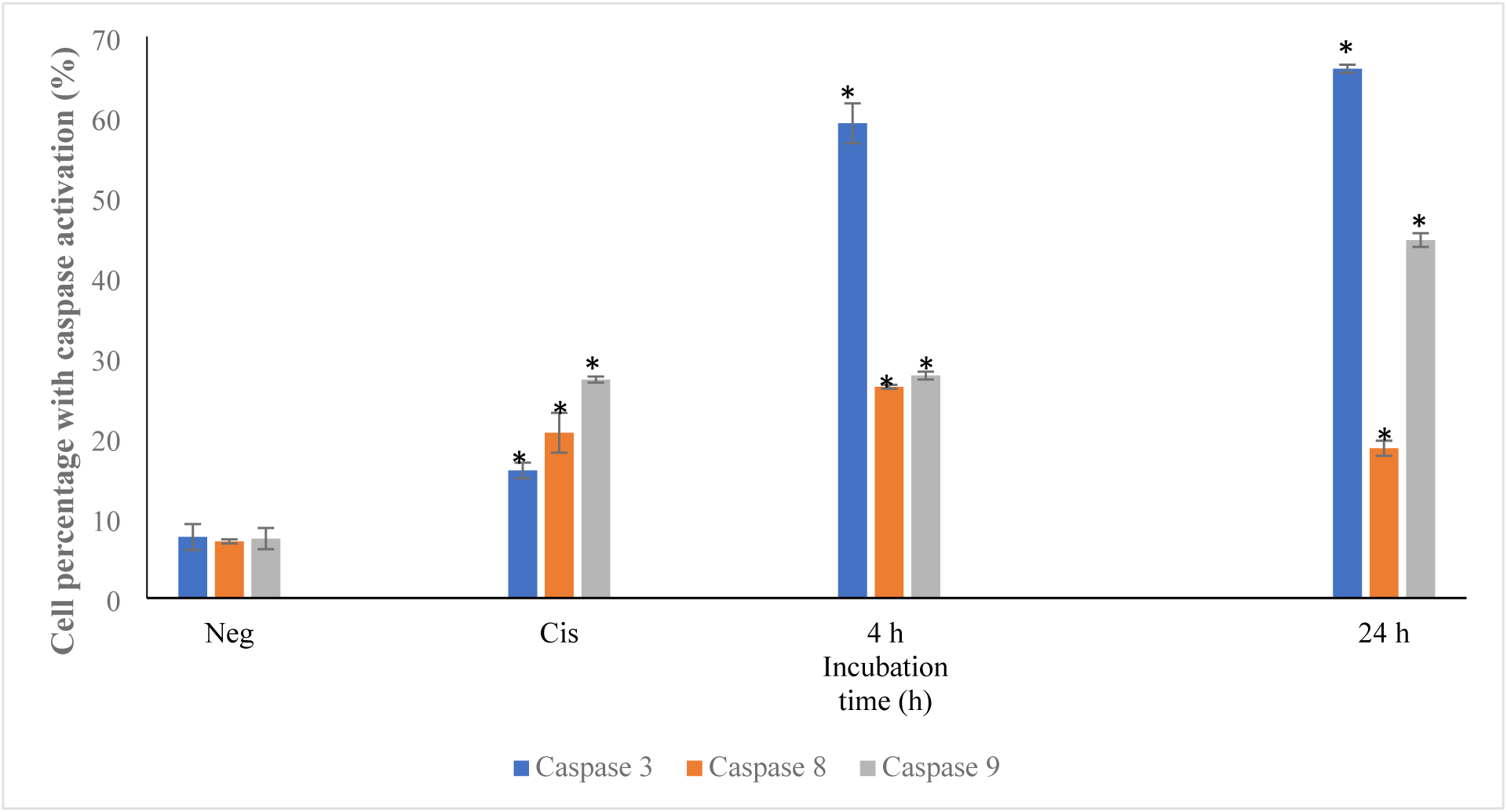
Caspase-8, -9, and -3 activity in A549 cells treated with DioSn-2 at IC₅₀ (1.86 μM). Data represent mean ± S.E.M from three independent experiments. \**p* < 0.05 vs. negative control. DISCUSSION & FUTURE WORK

### Synthesis and characterization of DioSn and TriSn

Four organotin(IV) *N*-alkyl-*N*-benzyldithiocarbamate compounds were successfully synthesized via an *in situ* method, adapted from Awang et al. [32] and Muthalib & Baba [33], with modifications including the use of 25% aqueous ammonia as a basic medium to facilitate the reaction and extend the reaction time, thereby improving the overall yield. The compounds were characterized by elemental analysis (C, H, N, and S) and spectroscopic techniques, including Fourier Transform Infrared (FT-IR) and nuclear magnetic resonance (NMR) spectroscopy (δ¹H, δ¹³C, and δ¹¹⁹Sn). Elemental compositions were consistent with the proposed molecular formula, RSn[S₂CN(L)]₄₋, where L = CH₃(C₆H₅) for DioSn-**1** (n = 2, R = CH₃) and DioSn-**2** (n = 2, R = C₆H₅); and L = CH₃(C₆H₅) for TriSn-**3** (n = 3, R = C₆H₅) and L = CH₂CH₃(C₆H₅) for TriSn-**4** (n = 3, R = C₆H₅).

FT-IR analysis revealed two prominent absorption bands characteristic of dithiocarbamate ligands: the thioureide υ(C=N) band observed at 1479–1497 cm⁻¹ and the υ(C–S) band at 950–998 cm⁻¹. Additionally, the Sn–S stretching vibration was detected in the range of 369–388 cm⁻¹. These findings are consistent with previous reports by Brown et al. [34] and Nomura et al. [35], who observed the thioureide band within 1450–1550 cm⁻¹, indicative of a polar C=N⁺ bond or partial double-bond character. The observed υ(C–S) absorption also aligns with typical dithiocarbamate features reported in the 950–1050 cm⁻¹ range [14]. The Sn–S stretching frequency is in agreement with earlier studies, which reported values between 325 and 390 cm⁻¹ [25,36,37].

Three NMR spectra (δ¹H, δ¹³C, and δ¹¹⁹Sn) were recorded to confirm the structural features of the compounds. In the δ¹H NMR spectrum, methylene protons attached to the phenyl ring appeared as singlets at δ 5.060–5.255 ppm [15]. The δ¹³C NMR spectra showed signals at δ 197–201 ppm, attributed to the thioureide carbon (–NCS₂), indicating C=N double bond character [38]. The δ¹¹⁹Sn NMR spectra for compounds 1–4 displayed chemical shifts from –497 to –180 ppm, consistent with coordination to Sn(IV) centers.

Single-crystal X-ray diffraction analysis of DioSn-**1** and TriSn-**3** confirmed the anisobidentate coordination mode of the dithiocarbamate ligands to Sn(IV), resulting in distorted geometries such as skewed octahedral (DioSn-**1**) and distorted trigonal bipyramidal (TriSn-**3**). In both compounds, the ligands coordinated through the Sn1–S2 bond. The Sn1–S1bond lengths in DioSn-**1** and TriSn-**3** were measured at 2.5381(6) Å and 2.4724(4) Å, respectively, and were notably shorter than the corresponding Sn1–S2 bond lengths (2.982(6)–3.020(4) Å), indicating that the Sn1–S2 interaction is weak and does not form a strong covalent bond. The coordination geometry of DioSn-1 can be described as skewed octahedral, typical of R₂Sn(S₂CNR’R”)₂complexes, based on the C1–Sn1–C1A arrangement [39]. In contrast, the molecular structure of TriSn-**3** exhibits a distorted trigonal bipyramidal geometry with anisobidentate ligand coordination, as inferred from the S–Sn–C and C–Sn–C bond angles. The anisobidentate coordination leads to unequal Sn–S bond lengths, resulting in structural asymmetry, consistent with earlier findings by Kana et al. [40]. Additionally, a torsion angle of Sn–S–C–S = 11.23(9)° indicates a slightly planar arrangement of the dithiocarbamate moiety. Overall, DioSn-**1** is six-coordinate, while TriSn-**3** is five-coordinate. The crystal packing of DioSn-**1** and TriSn-**3** is illustrated in Figures 1 and 2, respectively, while the C—H···π interactions in the crystal packing of TriSn-**3** are summarized in Table 7. No classical hydrogen bonding was observed in either compound; however, C—H···π interactions play a role in stabilizing the crystal structure of TriSn-**3**. The Donor—H···π centroid angles range between 138° and 165°, suggesting relatively strong interactions within the crystal packing.

**Table 7:** Geometry of C—H···π Interactions (Å,°) in the Crystal Packing of TriSn-3.

| **D—H⋯A** | **D—H** | **H⋯A** | **D⋯A** | **D—H⋯A (°)** |
| --- | --- | --- | --- | --- |
| C5—H5A⋯Cg5<sup>(i)</sup> | 0.9300 | 2.92 | 3.8217(16) | 165 |
| C12—H12A⋯Cg6<sup>(ii)</sup> | 0.9300 | 2.97 | 3.7604(18) | 142 |
| C23—H23A⋯Cg7<sup>(iii)</sup> | 0.9300 | 2.87 | 3.6107(16) | 138 |

(i) 1-x, 1-y, 1-z, (ii) 2-x, 1-y, 1-z, (iii) 1+x, y, z. Cg5 = C10–C15, Cg6 = C1–C6, Cg7 = C16–C21

**Table 8:**
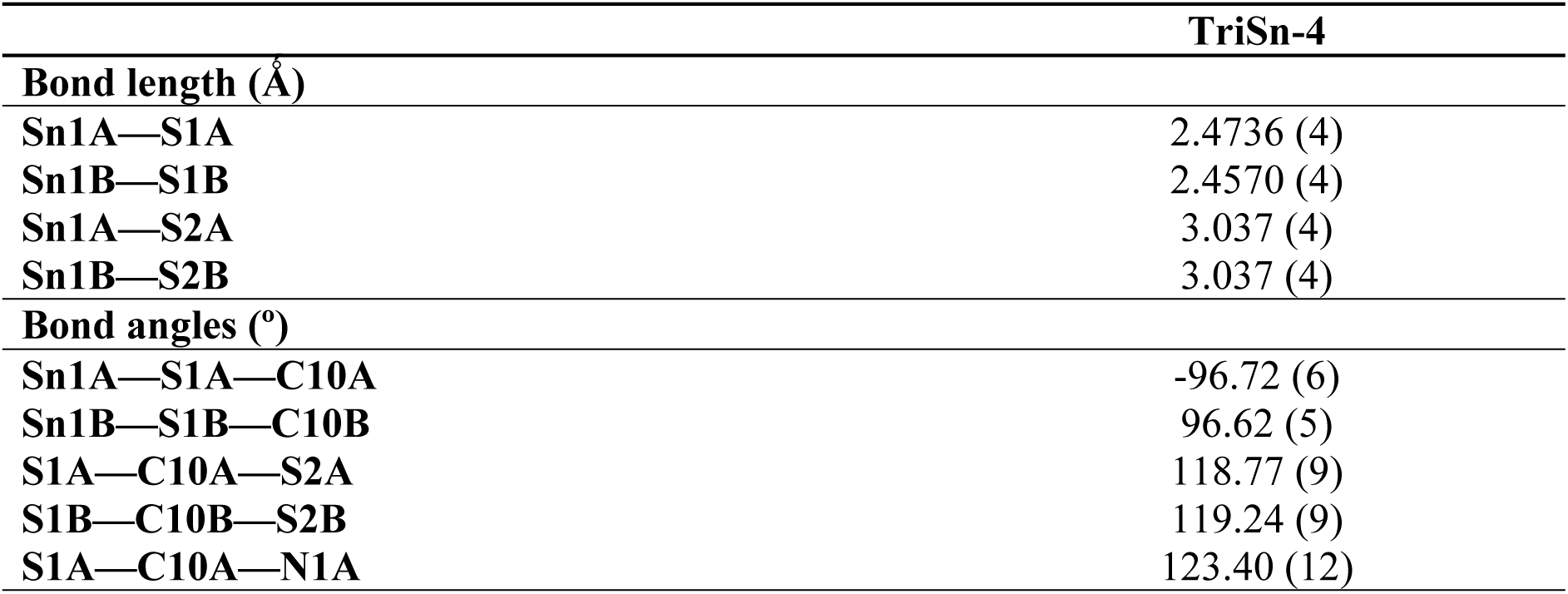
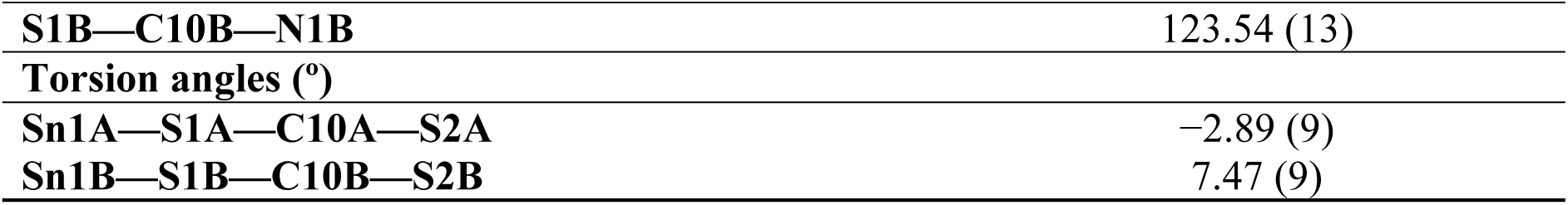
The selected geometric parameters of TriSn-4.

**Table 9:** IC_50_ and Selectivity Index of DioSn and TriSn Compounds in A549 and MRC-5.

| Compound | IC<sub>50</sub> (μM) ± SEM | | Selectivity index (SI) |
| --- | --- | --- | --- |
|  | A549 | MRC-5 |  |
| **Cisplatin (Positive Control)** | 32.00 ± 1.53 | 11.60 ± 0.31 | 0.36 |
| **1** | > 50.00 | - | - |
| **2** | 1.86 ± 0.15 | 12.00 ± 1.59 | 6.45 |
| **3** | 0.57 ± 0.13 | 0.64 ± 0.13 | 1.12 |
| **4** | 0.52 ± 0.06 | 0.70 ± 0.16 | 1.35 |

For comparison, the crystal structure of TriSn-**4**, previously reported by Abd Aziz et al. [21], also exhibited a five-coordinate distorted trigonal bipyramidal geometry, similar to TriSn-**3**. This suggests that organotin(IV) compounds with similar structural motifs tend to adopt comparable coordination geometries, despite variations in the dithiocarbamate ligands. In the case of DioSn-**2**, attempts to crystallize the compound resulted in poor-quality single crystals, likely due to structural instability in solution or the formation of microcrystals unsuitable for X-ray diffraction analysis [41].

### MTT cytotoxicity screening of DioSn and TriSn

The MTT assay revealed that compounds **2**–**4** exhibited significantly (p < 0.05) lower IC₅₀ values (0.52–1.86 μM) compared to cisplatin (32 μM), as shown in Figures 4–6. These findings align with previous studies where organotin(IV) compounds, such as Ph₃Sn(Boc-Orn-O), demonstrated 20–60 times higher antiproliferative activity than cisplatin across various cancer cell lines [42]. Similar results have been reported for organotin(IV) derivatives, which exhibit strong anticancer effects at low doses against several cancer types, including ovarian, lung, epidermoid, lymphoma, and cervical cancers [43]. In contrast, DioSn-**1** showed no significant cytotoxicity after 24 hours, even at 50 μM (27.07 μg/mL), indicating low toxicity (IC₅₀ > 25 μg/mL) [44]. These results suggest that cytotoxic activity is influenced by the compound’s structure, particularly the nature and number of alkyl groups bonded to the tin(IV) center [14]. The enhanced activity of compounds **2**–**4** may be attributed to the presence of phenyl groups on the tin atom and the aromatic ring in the dithiocarbamate ligand, which likely promote π-π interactions with biomolecules. Organotin compounds with phenyl substituents often display higher cytotoxicity than those bearing butyl or methyl groups [45,46].

Notably, compounds **2** to **4** exhibited selective cytotoxicity toward cancer cells (SI > 1) [47,48,49]. DioSn-**2**, in particular, showed potent cytotoxicity against A549 cells with an IC₅₀ of 1.86 μM, demonstrating approximately six-fold selectivity over normal human lung fibroblasts (MRC-5). As highlighted by Subramani et al. [50], a high selectivity index indicates a safer and more effective anticancer agent, characterized by minimal toxicity to normal cells while retaining strong activity against cancer cells. The role of ligands is also critical; coordination of ligands to the Sn(IV) center significantly enhances cytotoxic activity compared to the free ligands [51,52] supporting the hypothesis of a synergistic effect between the ligand and organotin moiety [53]. Chelation facilitates targeted delivery by stabilizing the compound and minimizing undesirable interactions with biomolecules, while the organotin(IV) fragment contributes directly to cytotoxicity [14]. Thus, the enhanced and selective cytotoxic effects observed in these compounds, compared to cisplatin, may result from the unique coordination and functional properties of the dithiocarbamate ligands.

### Apoptotic Morphology of A549 Cells Induced by DioSn-2, TriSn-3, and TriSn-4

Apoptosis is a key target in anticancer drug development. Compounds that trigger this pathway, particularly through modulation of proteins like the Bcl-2 family, hold strong potential as targeted therapies [54,55]. Unlike necrosis, apoptosis does not elicit inflammation, making it a more desirable mechanism for cancer treatment [56]. In this study, Annexin V-FITC/PI staining revealed that DioSn-**2**, TriSn-**3**, and TriSn-**4** induced apoptosis in over 50% of A549 cells—surpassing the effect of cisplatin—while necrotic cell levels remained low (1.70–3.73%) (Figure 9). These findings align with previous studies on organotin(IV) dithiocarbamates [14,15,48,49]. Additionally, apoptotic features in A549 cells treated with these compounds at IC₅₀ concentrations were observed microscopically, as shown in Figure 8.

For TriSn-**4**, over 80% apoptosis was observed, contrasting with the ∼50% cell viability measured by the MTT assay—a discrepancy also reported by Rasli et al. [48]. This difference stems from the distinct principles of each assay. MTT assesses metabolic activity via mitochondrial enzymes, while the Annexin V-FITC/PI assay uses flow cytometry to quantify live, early apoptotic, late apoptotic, and necrotic cells. Annexin V binds to phosphatidylserine exposed on apoptotic cell membranes, whereas propidium iodide (PI) stains DNA in cells with compromised membranes, marking late apoptosis or necrosis [57]. Notably, MTT cannot distinguish between reduced metabolism (cytostatic effects) and actual cell death (cytotoxic effects), which must be clearly differentiated [58]. Thus, combining assays is crucial for a comprehensive assessment of compound-induced cytotoxicity.

### DioSn-2 selectively induces cytotoxicity and mitochondria-mediated apoptosis in A549 cells

DioSn-**2** was selected for mechanistic studies due to its distinct chemical features and strong cytotoxic activity. FT-IR analysis confirmed the presence of Sn–S bonds, indicating bidentate coordination of the *N*-methyl-*N*-benzyldithiocarbamate ligand and suggesting structural stability [51,59]. ^13^C NMR spectra confirmed the presence of phenyl groups, which are known to enhance cytotoxicity through increased π–π interactions and lipophilicity [15,38, 53]. Additionally, phenyl substitution has been associated with reduced polarity and dipole moment around the tin atom, potentially facilitating improved membrane permeability [60,61]. Biologically, DioSn-**2** exhibited a six-fold lower IC₅₀ against A549 lung cancer cells than against normal human lung fibroblasts (MRC-5), indicating a favorable selectivity index [50]. Annexin V-FITC/PI staining showed that DioSn-**2** significantly (p < 0.05) induced apoptosis in ∼60% of A549 cells, with 5% early and 54% late apoptotic events. In contrast, cisplatin induced 14% early and 34% late apoptosis. The dominance of late apoptosis highlights DioSn-**2**’s potent cytotoxic effect, which exceeds the primarily cytostatic response seen with cisplatin [58]. These findings support DioSn-**2**’s selective anticancer activity via mitochondria-mediated apoptosis, particularly in rapidly proliferating cancer cells [62,63], positioning it as a promising candidate for further drug development.

The alkaline comet assay revealed that DioSn-**2** induces DNA damage as early as 30 minutes post-treatment, with damage severity increasing over time. This is supported by nuclear imaging, which showed progressively longer comet tails with extended treatment duration (Figure 10A). At all time points, DioSn-**2** caused significantly higher DNA tail moments compared to the negative control (0.52 ± 0.03, *p* < 0.05) and even exceeded the positive control (1.06 ± 0.02). These findings align with Syed Annuar et al. [49], who reported early DNA damage in K562 leukemia cells treated with Ph₃Sn(*N*,*N*-diisopropyldithiocarbamate) (OC2). DNA damage can trigger various cellular responses depending on severity—ranging from cell cycle arrest to apoptosis or necrosis [64,65]. Organotin(IV) compounds exert their genotoxic effects through different mechanisms. Some inhibit DNA synthesis by blocking nuclear protein synthesis [53,66], while phenyl-substituted derivatives can intercalate into DNA due to their planar structures, disrupting its integrity [67,68]. Consequently, di- and triphenyltin(IV) compounds exhibit stronger cytotoxic activity compared to their butyl or methyl analogs [46].

Organotin(IV) compounds are known to disrupt cytoskeletal and mitochondrial function, inducing apoptosis via mitochondrial depolarization, ROS generation, and loss of membrane potential (ΔΨm) [66,69]. In this study, treatment of A549 cells with DioSn-**2** at its IC₅₀ concentration resulted in a significant reduction in ΔΨm (14.2 ± 0.51%) within 1 hour, as detected by TMRE staining (p < 0.05). Even slight decreases in ΔΨm can impair ATP production and increase ROS [70], and the inherently higher ΔΨm in cancer cells makes mitochondria a selective therapeutic target [71,72]. Consistent with this, DioSn-**2** also induced time-dependent ROS generation, measured by DHE staining. Although early increases at 30 minutes were not statistically significant, ROS levels rose progressively, reaching a significant increase at 4 hours (29.6 ± 2.72%; p < 0.05). These findings align with previous studies where organotin(IV) compounds caused mitochondrial dysfunction and ROS-mediated apoptosis [49,73].

While moderate ROS levels support tumor growth, excessive ROS surpassing the cytotoxic threshold induces apoptosis, making mitochondrial ROS a promising anti-cancer target [74,75]. To further confirm ROS involvement in DioSn-**2**-induced apoptosis, A549 cells were pretreated with the antioxidant *N*-acetyl-*L*-cysteine (NAC) for 1 hour before DioSn-**2** exposure. NAC is known to neutralize oxidative stress and inhibit ROS-mediated apoptosis [76]. NAC (5 mM) pre-treatment significantly reduced apoptosis in A549 cells treated with DioSn-**2** (from 59.90% to 7.43%, *p* < 0.05), confirming that ROS plays a key role in apoptosis induction. This is consistent with findings by Celesia et al. [77] and Syed Annuar et al. [49], who showed that NAC reduces ROS and apoptosis caused by organotin compounds in colon and leukemia cells, respectively.

Interestingly, this study suggests a distinct mechanism for DioSn-**2**, where DNA damage precedes mitochondrial depolarization and ROS production, indicating early genotoxic stress as a trigger for apoptosis. This sequence—DNA damage (30 min), ΔΨm loss (1 h), and ROS increase (4 h)—supports a multi-step apoptotic pathway. While organotin(IV) compounds typically induce apoptosis through excessive ROS generation, mitochondrial membrane potential (ΔΨm) depolarization, and nuclear damage [74,75], our findings revealed that DioSn-**2** initiates apoptosis via early DNA damage, followed by ΔΨm reduction and delayed ROS production in A549 cells. These findings are consistent with those of Syed Annuar et al. [49], who reported that organotin(IV) dithiocarbamates can induce the intrinsic mitochondrial apoptotic pathway in response to DNA damage, an early apoptotic signal. This, in turn, leads to the loss of mitochondrial membrane potential (MMP) and the generation of reactive oxygen species (ROS), followed by increased activities of caspase-9, caspase-3, and poly(ADP-ribose) polymerase (PARP), thereby indicating that Ph₃Sn(*N*,*N*-diisopropyldithiocarbamate) induces intrinsic mitochondria-mediated apoptosis.

Mitochondria regulate apoptosis through components like the membranes, intermembrane space (IMS), and matrix [78]. Excess Ca²⁺ and oxidative stress trigger mitochondrial permeability transition pore (MPTP) opening, causing loss of membrane potential (ΔΨm), ATP depletion, and ROS overproduction [79,80]. This leads to the release of apoptotic proteins and mtDNA, triggering Bax/Bak activation, mitochondrial outer membrane permeabilization (MOMP), and caspase-mediated apoptosis [81]. Cytochrome c, normally part of the electron transport chain, becomes a pro-apoptotic signal when released into the cytosol. It binds Apaf-1, forming the apoptosome, which activates caspase-9 and downstream executioner caspases like caspase-3 and -7 [81,82]. In this study, DioSn-**2** significantly (p < 0.05) increased caspase-9 and -3 activity in A549 cells at 4 and 24 hours, with greater activation than caspase-8, suggesting intrinsic apoptosis. Cisplatin showed similar effects. Results from comet, TMRE, DHE, and caspase assays confirm that DioSn-**2** induces mitochondrial-mediated apoptosis via DNA damage, ΔΨm loss, and ROS generation, leading to caspase-9 and -3 activation. This aligns with previous findings where organotin compounds, such as tributyltin [83] and Ph_3_Sn(*N*,*N*-diisopropyldi- thiocarbamate) [49], triggered intrinsic apoptosis through mitochondrial disruption, oxidative stress, and apoptosome formation. The mechanism of action of DioSn-**2** on A549 cells at the IC₅₀ concentration is summarized in Figure 14.

**Figure 14.**
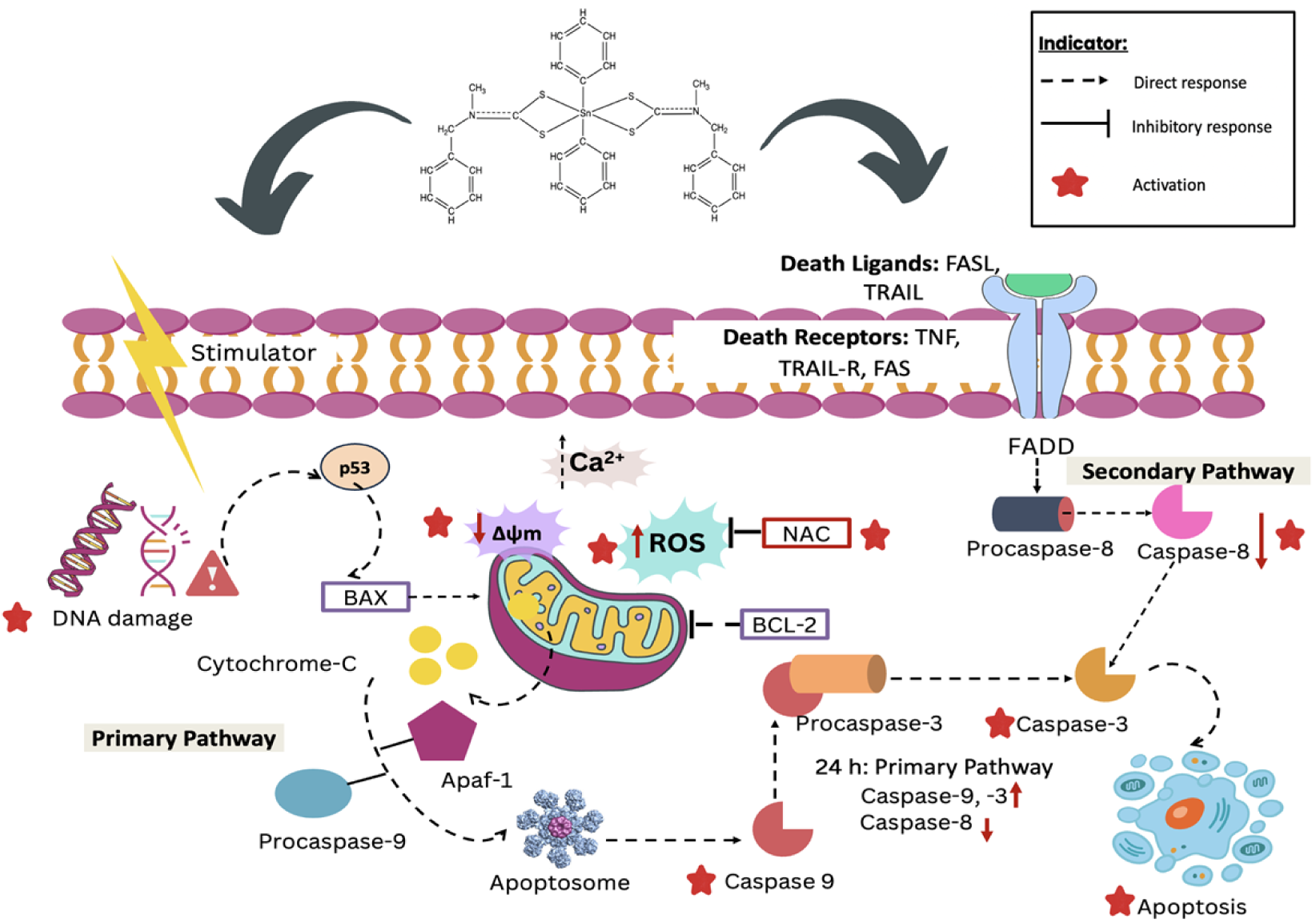
Proposed flowchart of the primary mechanism of action of DioSn-**2** on A549 cells at the IC₅₀ concentration.

## CONCLUSION

In conclusion, this study investigated the anticancer activity of four newly synthesized organotin(IV) dithiocarbamates—DioSn-**1**, DioSn-**2**, TriSn-**3**, and TriSn-**4**—in A549 lung cancer cells. Structural confirmation through elemental and spectroscopic analyses supported the proposed molecular compositions. Cytotoxicity assays revealed that DioSn-**2**, TriSn-**3**, and TriSn-**4** exhibited stronger cytotoxic effects than cisplatin, while DioSn-**1** was inactive. Apoptotic features such as cell shrinkage and membrane blebbing were observed in treated cells. Among them, DioSn-**2** selectively induced mitochondria-mediated apoptosis, marked by DNA damage, loss of mitochondrial membrane potential (Δψm), and excessive ROS generation. The role of oxidative stress was confirmed by a significant reduction (p < 0.05) in apoptosis upon NAC pre-treatment. These events led to the activation of caspase-9 and caspase-3, confirming the involvement of the intrinsic apoptotic pathway. Overall, our findings highlight the potent anti-proliferative effects of these novel compounds and their promise as metal-based chemotherapeutic agents. Further studies should explore their molecular mechanisms and evaluate their efficacy across various cancer models, paving the way for the development of next-generation organotin-based anticancer therapies.

## ACKNOWLEDGEMENT

The authors express their sincere appreciation to the Faculty of Health Sciences, Universiti Kebangsaan Malaysia (UKM), Sunway University, and the Center for Research and Instrumentation Management (i-CRIM), UKM, for providing access to essential laboratories and facilities that were crucial for the successful completion of this research.

## FUNDING

This work was supported by the Fundamental Research Grant Scheme (FRGS) [grant number: FRGS/1/2021/STG04/UKM/02/5] from the Ministry of Higher Education (MOHE), Malaysia.

## DATA AVAILABILITY STATEMENT

All data generated or analyzed during this study are included in this published article.

## AUTHOR CONTRIBUTIONS STATEMENT

N.A : Supervision, conceptualization, Methodology

N.A.A.A: Writing original manuscript, Revised original manuscript, Methodology

N.Z.Z: Writing original manuscript, Methodology

N.F.K: Supervision A.H: Supervision

N.N.M.A: Supervision

K.M.C: Supervision

## Notes

### Competing Interest Statement

The authors have declared no competing interest.

